# The asymmetric opening of HIV-1 Env by a potent CD4 mimetic enables anti-coreceptor binding site antibodies to mediate ADCC

**DOI:** 10.1101/2024.08.27.609961

**Authors:** Jonathan Richard, Michael W. Grunst, Ling Niu, Marco A. Díaz-Salinas, William D. Tolbert, Lorie Marchitto, Fei Zhou, Catherine Bourassa, Derek Yang, Ta Jung Chiu, Hung-Ching Chen, Mehdi Benlarbi, Guillaume-Beaudoin-Buissières, Suneetha Gottumukkala, Wenwei Li, Katrina Dionne, Étienne Bélanger, Debashree Chatterjee, Halima Medjahed, Wayne A. Hendrickson, Joseph Sodroski, Zabrina C. Lang, Abraham J. Morton, Rick K. Huang, Doreen Matthies, Amos B. Smith, Walther Mothes, James B. Munro, Marzena Pazgier, Andrés Finzi

## Abstract

HIV-1 envelope glycoproteins (Env) from primary HIV-1 isolates typically adopt a pretriggered “closed” conformation that resists to CD4-induced (CD4i) non-neutralizing antibodies (nnAbs) mediating antibody-dependent cellular cytotoxicity (ADCC). CD4-mimetic compounds (CD4mcs) “open-up” Env allowing binding of CD4i nnAbs, thereby sensitizing HIV-1-infected cells to ADCC. Two families of CD4i nnAbs, the anti-cluster A and anti-coreceptor binding site (CoRBS) Abs, are required to mediate ADCC in combination with the indane CD4mc BNM-III-170. Recently, new indoline CD4mcs with improved potency and breadth have been described. Here, we show that the lead indoline CD4mc, CJF-III-288, sensitizes HIV-1-infected cells to ADCC mediated by anti-CoRBS Abs alone, contributing to improved ADCC activity. Structural and conformational analyses reveal that CJF-III-288, in combination with anti-CoRBS Abs, potently stabilizes an asymmetric “open” State-3 Env conformation, This Env conformation orients the anti-CoRBS Ab to improve ADCC activity and therapeutic potential.

## MAIN

Combination antiretroviral therapy (cART) can inhibit multiple stages of the HIV-1 life cycle, suppressing viral replication and preventing the progression to AIDS in people living with HIV-1 (PLWH)^1,2^. However, cART alone does not eradicate the virus. Due to the persistence of a latent reservoir, lifelong cART is required to prevent viral rebound^3–6^. New strategies aimed at targeting and eliminating HIV-1-infected cells are needed to achieve a cure. Antibody-dependent cellular cytotoxicity (ADCC) represents one of the primary effector mechanisms used by the immune system to clear infected cells. This response relies on the ability of antibodies (Abs) to bind viral antigens on the surface of infected cells, thereby facilitating their recognition and destruction by immune effector cells such as natural killer (NK) cells. The HIV-1 envelope glycoprotein (Env) trimer represents the only viral protein exposed on the surface of HIV-1-infected cells and therefore serves as the primary target for ADCC-mediating antibodies^7,8^.

Env is derived from a gp160 precursor, which is synthesized, trimerized and glycosylated in the host cell’s endoplasmic reticulum^7,9^. Trimeric Env is subsequently cleaved by host furin-like proteases as it traffics through the Golgi apparatus on its way to the cell surface^10–12^. The cleaved mature Env trimer is composed of three exterior gp120 subunits non-covalently associated with three transmembrane gp41 subunits^13–15^. This trimeric Env is dynamic, transitioning from a pretriggered “closed” (State 1) to a “fully open” (State 3) conformation upon interaction with the viral receptor CD4^16–18^. During viral entry, CD4 engages the gp120 subunit by inserting phenylalanine 43 (Phe43) of domain 1 into a highly conserved pocket located at the interface of the inner and outer gp120 domains, known as the Phe43 cavity^19^. CD4 interaction stabilizes downstream Env conformations, leading to a progression from State 1, through an intermediate (State 2), to the fully open State-3 conformation^16,17,20^.

Unliganded Envs from most primary HIV-1 isolates naturally assume a “closed” State-1 conformation^16^, which is resistant to non-neutralizing antibodies (nnAbs) commonly elicited in the majority of PLWH^21,22^. Although these nnAbs have the potential to eliminate infected cells via ADCC, they target epitopes that only become accessible in “open” conformations, upon Env-CD4 interaction^22,23^. HIV-1 has evolved Nef and Vpu proteins that limit this possibility by downregulating CD4, thereby protecting HIV-1-infected cells from ADCC mediated by CD4-induced (CD4i) nnAbs^22–24^. By driving Env to downstream “open” conformation(s), CD4-mimetic compounds (CD4mcs) expose vulnerable CD4i Env epitopes and sensitize HIV-1-infected cells to ADCC mediated by Abs present in the plasma from PLWH^25–30^.

Two families of CD4i nnAbs, the anti-cluster A and anti-coreceptor binding site (CoRBS) Abs, have been shown to mediate potent ADCC when combined with BNM-III-170, a lead CD4mc compound of the indane class^31,32^. This sensitization of infected cells to ADCC requires a sequential opening of the Env trimer^32^. In contrast to membrane-bound CD4, BNM-III-170 is unable on its own to fully “open-up” the trimer-^31^. However, BNM-III-170 efficiently exposes the CoRBS and subsequent binding of anti-CoRBS Abs further opens the Env trimer, exposing the inner domain of gp120 and enabling the binding of anti-cluster A Abs^32^. Of note, it is well described in the literature that anti-CoRBS, on their own, poorly mediate ADCC^32–36^.

Recently, more potent CD4mcs have been developed^26^. These new CD4mcs, based on an indoline scaffold, exhibit increased potency and breadth in preventing viral entry over the previous lead indane CD4mc, BNM-III-170. This enhanced antiviral activity is likely due to more favorable π–π overlap from the indoline pose and better contacts with the vestibule of the CD4-binding pocket on gp120^26^. The lead indoline CD4mc, CJF-III-288, also potently sensitizes HIV-1-infected cells to ADCC mediated by plasma from PLWH^25^. In the present study, we aimed to better understand the mechanism behind this improved activity. Unlike previous indane CD4mcs, we found that CJF-III-288 enables anti-CoRBS Abs to mediate ADCC in absence of anti-cluster A Abs, contributing to its improved ADCC activity. Mechanistically, we found that CJF-III-288 binding is associated with the stabilization of an asymmetric open State-3 Env conformation and a modification of anti-CoRBS Ab binding orientation.

## RESULTS

### CJF-III-288 enables anti-CoRBS Abs to mediate ADCC

To better characterize the ADCC-stimulating potential of the lead indoline CD4mc CJF-III-288, we compared its activity in combination with an anti-cluster A/anti-CoRBS Ab cocktail to that of the previous indane CD4mc BNM-III-170^32^. Activated human primary CD4+ T cells were infected with the transmitted/founder virus CH058 (CH058TF). Two days post-infection, the ability of anti-cluster A Abs (A32 or N5i5) and anti-CoRBS Ab (17b), to recognize and eliminate the HIV-1-infected cells by ADCC, alone or in combination, was evaluated by flow cytometry. As previously reported^32,33^, BNM-III-170 failed to expose the cluster A epitope on its own, but significantly improved 17b binding (Fig. 1a). Since 17b engagement exposes the cluster A epitope, more antibody binding was observed upon anti-cluster A addition. Despite efficiently recognizing HIV-1-infected cells in the presence of BNM-III-170, 17b failed to mediate ADCC alone (Fig. 1b). Elimination of infected cells by ADCC was only detected when 17b and an anti-cluster A Ab were combined together with BNM-III-170 (Fig. 1b). Intriguingly, despite similar levels of 17b binding upon CJF-III-288 addition, this antibody was able to mediate ADCC in the presence of this CD4mc alone (Fig. 1b). As observed with BNM-III-170, CJF-III-288 and 17b addition enabled anti-cluster A Ab interaction (Fig. 1a and Extended Data Fig. 1). Importantly, these phenotypes were consistent for three additional infectious molecular clones (IMC) (HIV-1_JRFL_, HIV-1_AD8_ and HIV-1_JRCSF_) (Fig. 1c,d) and for different anti-CoRBS Abs (N12i2, X5, C2, 412D, and 48D) (Fig. 1e,f). The capacity to sensitize HIV-1-infected cells to ADCC mediated by anti-CoRBS Abs was specific to the indoline CD4mc CJF-III-288 and was not observed with the indane (BNM-III-170) or the piperidine (TFH-I-116-D1) CD4mc^37^ (Fig.1e,f). These results indicate that the indoline CD4mc CJF-III-288 allows anti-CoRBS Abs to mediate ADCC against HIV-1-infected cells.

**Fig. 1.**
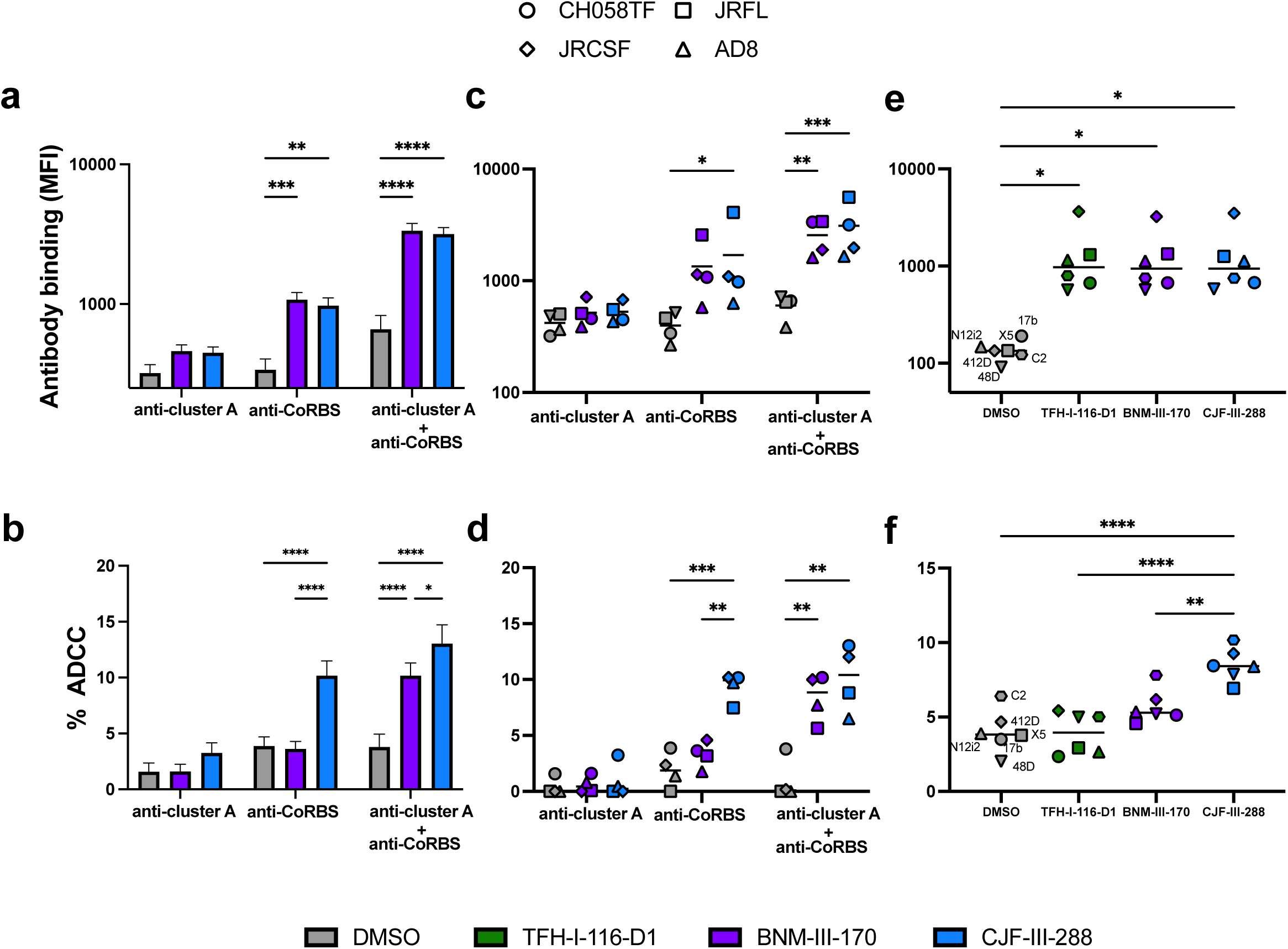
CJF-III-288 sensitizes HIV-1-infected cells to ADCC mediated by various anti-CoRBS Abs. **a**, Recognition and **b**, ADCC-mediated elimination of primary CD4+ T cells infected with HIV-1_CH058TF_ by anti-cluster A Abs (A32 or N5i5) and anti-CoRBS Abs (17b), alone or in combination (at 5µg/ml total concentration), in the presence of 50 µM of indicated CD4mc or equivalent volume of DMSO. **c**, Recognition and **d**, ADCC-mediated elimination of primary CD4+ T cells infected with indicated IMCs by anti-cluster A and anti-CoRBS, alone or in combination, in the presence of 50 µM of indicated CD4mc or equivalent volume of DMSO. **c-d**, Shown are the median fluorescence intensities (MFI) and percentage of ADCC obtained in at least 3 experiments for each IMC. **e,** Recognition and **f**, ADCC-mediated elimination of primary CD4+ T cells infected with HIV-1_CH058TF_ by indicated anti-CoRBS Abs in the presence of 50 µM of indicated CD4mc or equivalent volume of DMSO. **e-f**, Shown are the mean MFI and percentage of ADCC obtained from at least 3 experiments for each tested anti-CoRBS Abs (17b, N12i2, X5, C2, 412D and 48D). Statistical significance was tested using **a-b**, Two-way ANOVA test with a Tukey post-test, **c-f** one-way ANOVA with a Tukey post-test or Kruskal-Wallis test with a with a Holm-Sidak post-test based on the normality test. (* p<0.05,** p<0.01,*** p<0.001,**** p<0.0001)

### CJF-III-288 shows improved ADCC activity at low concentrations

We next compared the ability of CJF-III-288 and BNM-III-170 to induce recognition of infected cells and ADCC by anti-CoRBS Abs +/− anti-Cluster A Abs over a range of concentrations. Activated human primary CD4+ T cells were infected with the same four primary IMCs, and the ability of 17b and A32, alone or in combination, to recognize HIV-1-infected cells was assessed two days post-infection in the presence of increasing concentrations of each CD4mc. As presented in Fig. 2, both CD4mc similarly enhanced the ability of 17b alone or combined with A32, to recognize HIV-1-infected cells at the highest concentration tested (50µM for JRFL, JRCSF and AD8). Of note, since CH058 is more susceptible to CD4mc due to the presence of a threonine at position 375^38^, similar levels of binding were detected starting at 1 µM. At low concentrations (<10 µM for JRFL, JRCSF and AD8 and <1 µM for CH058), CJF-III-288 was more effective than BNM-III-170 (Fig. 2a,c) in enabling Abs to recognize infected cells. This improved potency was reflected in a significant increase in the area under the curve (AUC) of Ab binding for all tested IMCs (Fig. 2b,d). Supporting its improved potency, CJF-III-288 also exposed the CoRBS on Env from the clade A HIV-1_BG505_ virus, which was reported to be resistant to BNM-III-170^26^ (Extended Data Fig. 2). The improved potency of CJF-III-288 was also reflected in its capacity to sensitize infected cells to ADCC mediated by anti-CoRBS Abs alone or in combination with anti-Cluster A Abs. (Fig. 2e,g). This superior ADCC activity of CJF-III-288 against HIV-1_CH058TF_-infected cells was again reflected by a significant increase in the AUC (Fig. 2f,h). As expected, minimal recognition and little ADCC-mediated elimination of infected cells were observed with A32 in the absence of 17b (Extended Data Fig. 3).

**Fig. 2.**
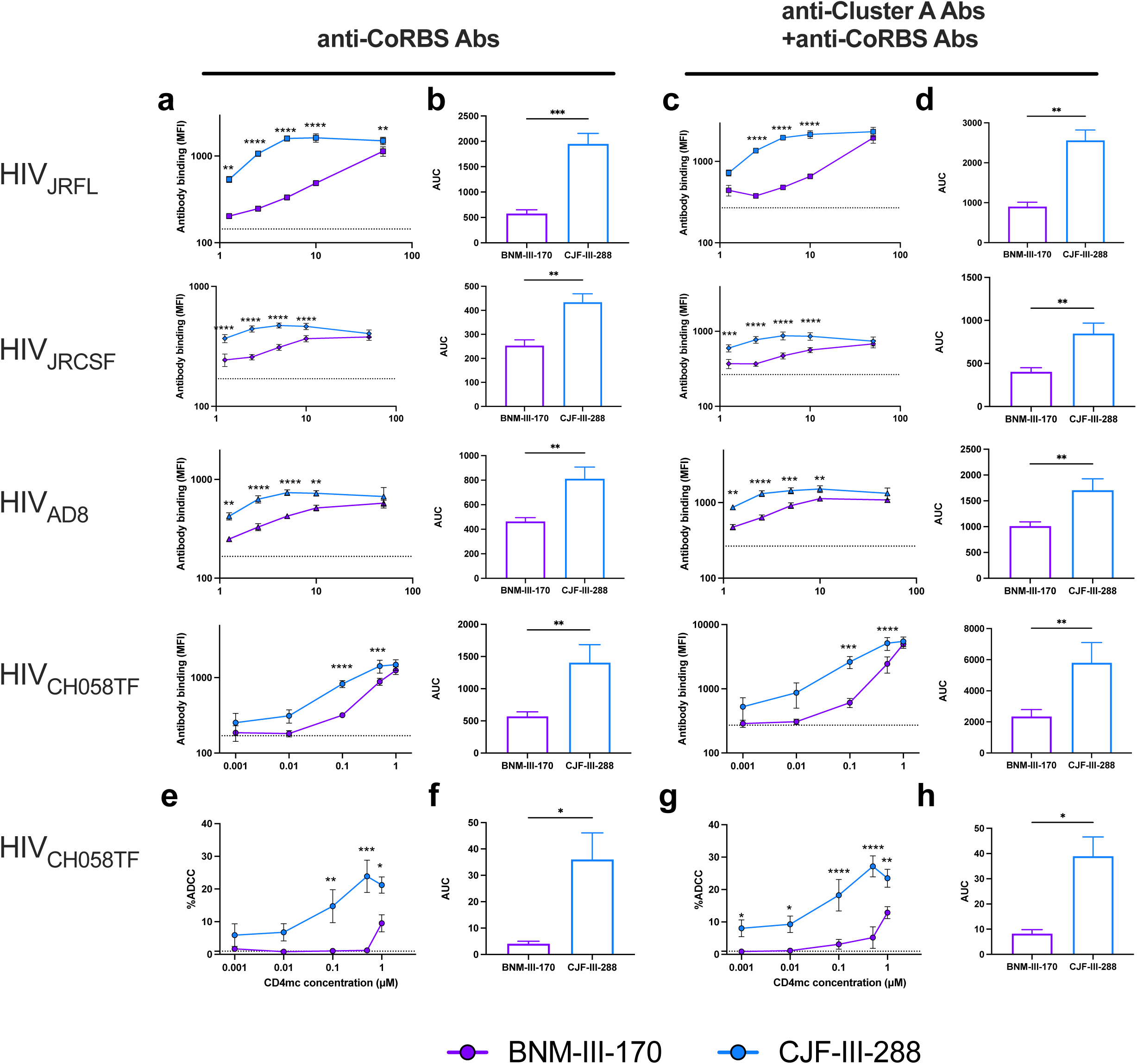
CJF-III-288 outperforms BNM-III-170 at lower concentrations. **a-d**, Recognition of primary CD4+ T cells infected with indicated IMC by anti-CoRBS Ab (17b) alone or in combination with anti-cluster A Ab (A32), in the presence of indicated concentration of BNM-III-170 or CJF-III-288. **a,c**, Graphs represent the median fluorescence intensities (MFI) obtained in at least 4 independent experiments with each IMCs. **b,d**, Graphs represent the area under the curve (AUC) calculated from the MFI obtained in **a** and **c**. **e-h**, ADCC-mediated elimination of CD4+ T cells infected HIV-1_CH058TF_ by anti-CoRBS Abs alone or in combination with anti-cluster A Abs, in the presence of indicated concentration of BNM-III-170 or CJF-III-288. **e,g**, Shown are percentage of ADCC obtained in at least 4 experiments. **f,h**, Shown are the area under the curve (AUC) calculated from the percentage of ADCC obtained in **e** and **g.** The dashed lines represent the mean binding or ADCC obtained in absence of CD4mc. Statistical significance was tested using **a,c,e,g**, Two-way ANOVA test with a Holm-Sidak post-test, **b,d,f,h** paired t test or Wilcoxon test based on the normality test. (* p<0.05,** p<0.01,*** p<0.001,**** p<0.0001).

To confirm these findings with primary clinical samples, CD4+ T cells were expanded from PLWH under ART, as previously described ^27,31,39^. Viral replication was monitored over time by intracellular p24 staining. Upon expansion, CD4+ T cells were stained with nnAbs in the presence of the indicated CD4mc. Expanded endogenously-infected cells were also used as target cells and autologous peripheral blood mononuclear cells (PBMCs) as effectors using a FACS-based ADCC assay. Consistent with the results obtained with IMC infection, superior antibody binding and ADCC responses were observed with CJF-III-288 relative to BNM-III-170 at different concentrations (Fig. 3).

**Fig. 3.**
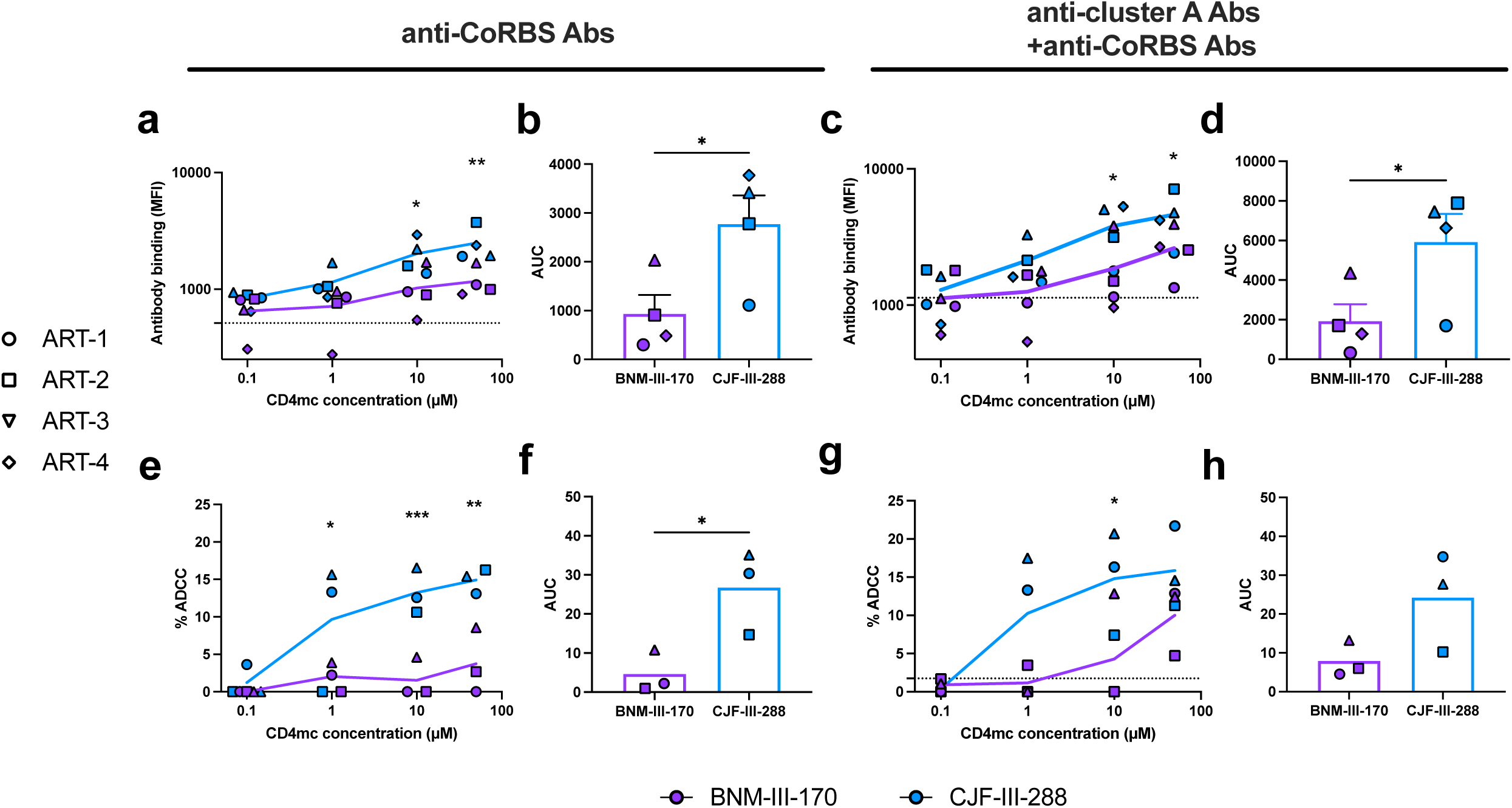
CJF-III-288 shows superior ADCC activity than BNM-III-170 against *ex-vivo-*expanded CD4+ T cells from PLWH. **a-d**, Recognition of *ex-vivo-*expanded CD4+ T cells from four PLWH under ART by anti-CoRBS Abs alone or in combination with anti-cluster A Abs, in the presence of indicated concentration of BNM-III-170 or CJF-III-288. **a,c**, Graphs represent MFI obtained with each donor. **b,d**, Graphs represent the AUC calculated from the MFI obtained in **a** and **c**. **e-h**, ADCC-mediated elimination of *ex-vivo-*expanded CD4+ T cells from three PLWH under ART by anti-CoRBS Abs alone or in combination with anti-cluster A Abs, in the presence of indicated concentration of BNM-III-170 or CJF-III-288. **e,g**, Shown are percentage of ADCC obtained with each donor. **f,h**, Shown are the area under the curve (AUC) calculated from the percentage of ADCC obtained in in **e** and **g**. The dashed lines represent the mean binding or ADCC obtained in absence of CD4mc. Statistical significance was tested using **a,c,e,g**, Two-way ANOVA test with a Holm-Sidak post-test, **b,d,f,h** paired t test or Wilcoxon test based on the normality test. (* p<0.05,** p<0.01,*** p<0.001,**** p<0.0001)

### Anti-CoRBS Abs drive PLWH plasma-mediated ADCC in the presence of CJF-III-288

We next extended our analysis to plasma from PLWH. As observed with nnAbs, CJF-III-288 was more effective than BNM-III-170 in enhancing plasma binding and ADCC activity at low concentrations (Fig. 4a-d). To evaluate the contribution of anti-CoRBS Abs to the improved ADCC activity mediated by PLWH plasma in the presence of CJF-III-288, we performed a Fab fragment competition assay. Briefly, HIV-1-infected cells were preincubated with 17b Fab fragments in the presence of CJF-III-288 prior to incubation with autologous PBMCs and PLWH plasma in the FACS-based ADCC assay. In this experimental setting, the 17b Fab masks the CoRBS exposed by CJF-III-288 and prevents the binding and ADCC activity of anti-CoRBS Abs present in PLWH plasma. Preincubation with 17b Fab substantially reduced the ADCC responses mediated by PLWH plasma in the presence of CJF-III-288, thus confirming the key contribution of anti-CoRBS Abs in this response (Fig. 4e).

**Fig. 4.**
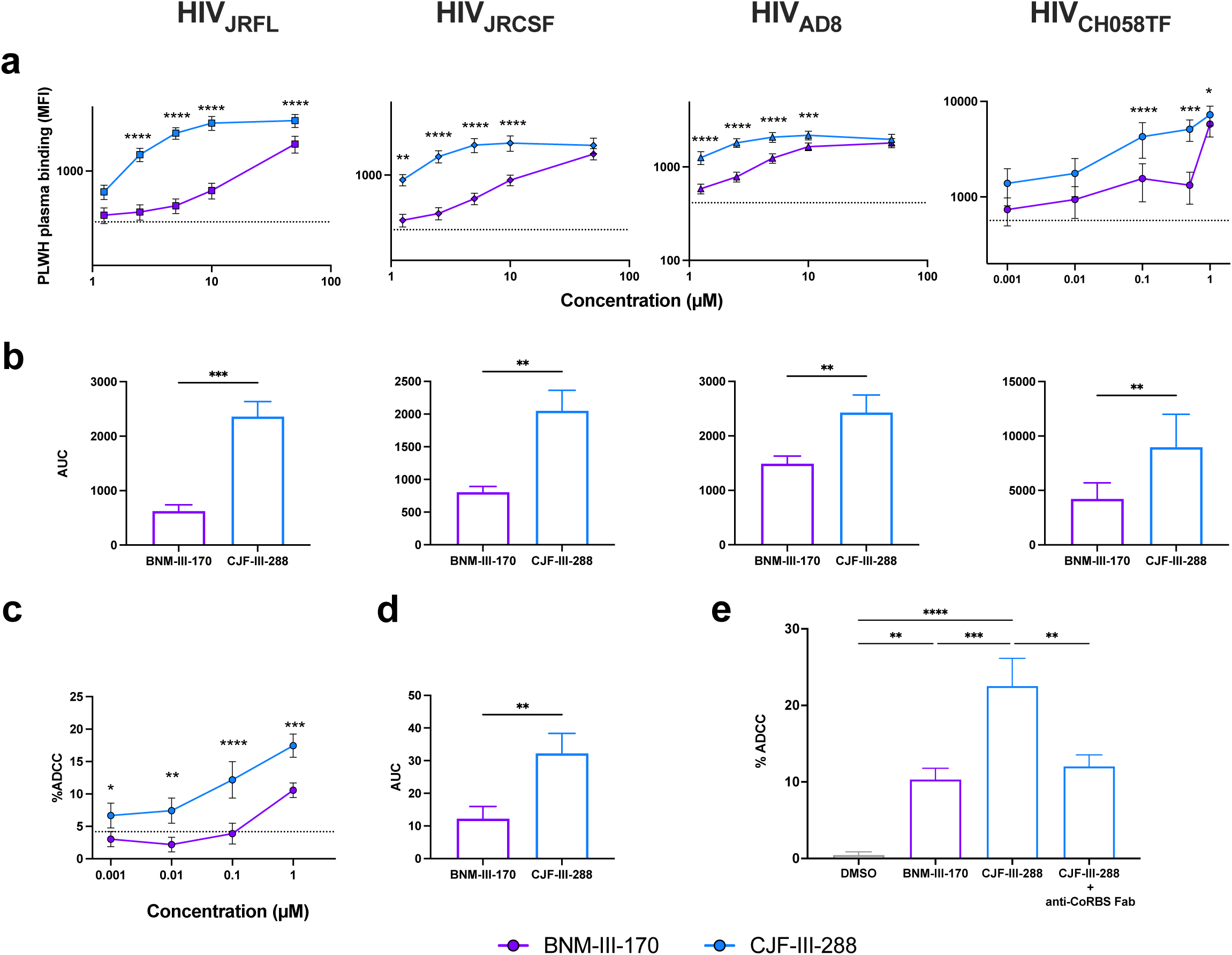
Anti-CoRBS Abs contribute to the improved ADCC activity of PLWH plasma in the presence of CJF-III-288. **a,b,** Recognition of primary CD4+ T cells infected with indicated IMC by PLWH plasma, in the presence of indicated concentration of BNM-III-170 or CJF-III-288. **a**, Shown are median fluorescence intensities obtained with plasma from 8 PLWH. **b**, Shown are the area under the curve (AUC) calculated from the MFI obtained in **a**. **c,d,** ADCC-mediated elimination of HIV-1_CH058TF_-infected primary CD4 T cells by plasma from PLWH in the presence of indicated concentration of BNM-III-170 or CJF-III-288. **c**, Shown are percentage of ADCC obtained with plasma from 8 PLWH. **d**, Shown are the AUC calculated from the ADCC values presented in **c**. The dashed lines represent the mean MFI or ADCC obtained in absence of CD4mc. **e**, ADCC-mediated elimination of HIV-1_CH058TF_-infected primary CD4 T cells preincubated or not with 17b Fab fragments, by plasma from PLWH in the presence of 1 µM of BNM-III-170 or CJF-III-288. Statistical significance was tested using **a,c**, Two-way ANOVA test with a Holm-Sidak post-test, **b,d,** paired t test or Wilcoxon test based on the normality test, **e**, One-way ANOVA with a Holm-Sidak post-test (* p<0.05,** p<0.01,*** p<0.001,**** p<0.0001)

### CJF-III-288, in combination with anti-CoRBS Abs, stabilizes the State-3 Env conformation

Our functional data indicated that CJF-III-288 could stabilize Env in a conformation distinct from that induced by other CD4mc, including BNM-III-170. To determine the impact of CJF-III-288 on Env conformation and how it differs from the one induced by BNM-III-170, we used a modified version of a well-established single-molecule Förster Resonance Energy Transfer (smFRET) imaging assay^16^. Briefly, one fluorophore was attached to the V1 loop of HIV-1_JRFL_ gp120 using amber stop codon suppression to introduce a non-natural amino acid (nnAA). The nnAA was then labelled by copper-free click chemistry with a tetrazine-conjugated fluorophore ^40^. A second fluorophore was attached to the A1 peptide inserted into the V4 loop of gp120^29^. Native virions incorporating a single labeled Env protomer were immobilized on quartz microscope slides and imaged using internal reflection fluorescence (TIRF) microscopy. As previously reported^16,17,29,31,41,42^, smFRET analysis of unbound Env showed a predominant low-FRET state, consistent with the adoption of a “closed” State-1 Env conformation (Fig. 5a,c). Incubation of the virions with the CD4mcs BNM-III-170 or CJF-III-288 led to destabilization of State 1 and a shift towards more open downstream conformations, including the high-FRET States 2 and 2A, and the intermediate-FRET State 3. This reduction of State 1 occupancy was significantly greater upon incubation with CJF-III-288 than with BNM-III-170 (Fig. 5c). Incubation with the anti-CoRBS Ab 17b alone had no effect on Env conformation. The combination of 17b and BNM-III-170 only modestly reduced State 1 occupancy, consistent with our previous results^31^. However, when combined with CJF-III-288, 17b significantly modified Env’s conformation, further reducing State 1 occupancy and favoring State 3 (Fig. 5a,c). Similarly, incubation of virions with CJF-III-288 in combination with PLWH plasma resulted in improved State 3 stabilization relative to BNM-III-170 (Fig. 5b,d). Stabilization of State 3, as well as State 1 destabilization by PLWH plasma correlated with ADCC (Fig. 5e). Altogether, these results indicate that CJF-III-288, in combination with anti-CoRBS Abs or plasma from PLWH, shifts the conformational landscape from State 1 towards the fully open State 3 conformation. Stabilization of the State-3 conformation could sensitize HIV-1-infected cells to ADCC mediated by anti-CoRBS Abs alone.

**Fig. 5.**
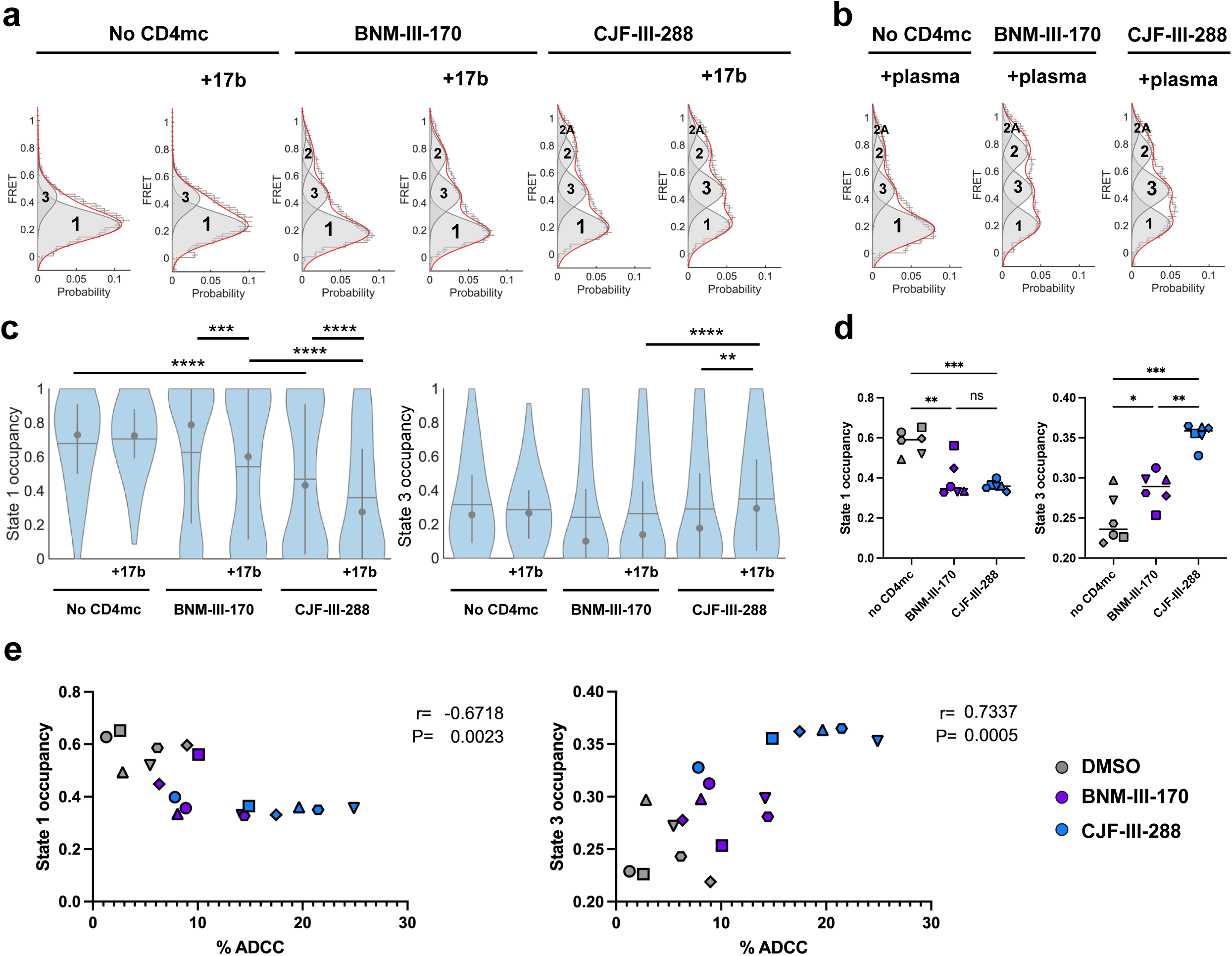
CJF-III-288 in combination with anti-CoRBS Abs stabilize Env State 3. **a,b** Histograms of FRET values compiled from the population of individual HIV-1_JRFL_Env trimers on the virion surface in the absence or presence of CD4mc (BNM-III-170 or CJF-III-288), the anti-CoRBS Ab 17b, or PLWH plasma. Histograms are presented as the mean determined from three technical replicates with error bars reflecting the standard error. Overlaid on the histograms are Gaussian distributions determined from HMM analysis of the individual FRET trajectories. Conformational state labels (States 1, 2, 2A, and 3), which have been previously identified ^16,31^, are indicated on the corresponding Gaussian. **c,d,** The occupancies in States 1 and State 3 were calculated from the HMM analysis for each trace and represented with violin plots. **c**, The mean and median occupancy are shown as horizontal lines and circles, respectively. Vertical lines reflect the 25-75% quantiles. e, Correlation between State 1 and State 3 occupancy and ADCC mediated by plasma from PLWH **d**,e, Each shape represents a different plasma from PLWH. Statistical significance was tested using (c,d) one-way ANOVA with a Holm-Sidak post-test and (e) Spearman rank correlation test (* p<0.05,** p<0.01,*** p<0.001,**** p<0.0001, ns: not significant).

### CJF-III-288 induces an asymmetric opening of Env that modifies the binding orientation of anti-CoRBS Abs

Previous studies have revealed that the ability of Abs to bind Fcγ receptors (FcγR) and activate ADCC is influenced by their binding orientation^36,43^. Based on this observation, we hypothesized that CJF-III-288 could stabilize Env in a conformation that modifies the orientation of anti-CoRBS Abs binding, thereby enhancing their ADCC activity. To gain molecular level insights into potential differences in Env trimer architecture, specifically the geometry of trimer assembly and the orientation of anti-CoRBS antibodies bound to Env triggered by CJF-III-288, we employed molecular imaging methods including single-particle Cryo-EM and CryoET. Using Cryo-EM, we determined a 3.5-Å structure of the BG505 sgp140 SOSIP.664 trimer in complex with CJF-III-288 and 17b Fab (Extended Data Table 1, Extended Data Figs. 4-5). We selected 17b as a representative anti-CoRBS Ab in the Cryo-EM studies to allow direct comparison to the complex of BNM-III-170-BG505 SOSIP.664-17b Fab determined previously using the same method by the Bjorkman laboratory^44^. Figure 6 shows the overall structure of the CJF-III-288-BG505 SOSIP.664-17b Fab complex that we determined using C1 symmetry. The overall density of gp120-gp41 protomers was well-defined with the some poorly resolved regions corresponding to the 17b Fab (Extended Data Fig. 4). There was high-resolution density within the Phe43 cavity in all gp120 protomers for CJF-III-288 (Extended Data Fig. 4d). The CJF-III-288-BG505 SOSIP.664-17b Fab trimer is an asymmetric assembly of the three protomers that differ mainly in the conformation of gp41 and the 17b Fab (only the variable portions of the Fabs were built into the model due to poor densities for the constant regions). Overall, the three CJF-III-288-BG505 SOSIP.664-17b Fab protomers can be superimposed with an average root-mean-square deviation (RMSD) for Cα atoms of 2.52 Å (the RMSD between protomer A and B, protomer A and C, and protomer B and C are 2.69 Å, 1.73 Å, and 3.14 Å respectively, Extended Data Fig. 5a). On the whole, the gp120 protomers are the best-defined portions of the complex with the conformation of residues forming the CJF-III-288 binding pocket within each protomer being well preserved (the RMSD between residues of the CJF-III-288 pockets of protomers A and B, protomers A and C, and protomer B and C are 0.565 Å, 0.563 Å, and 0.524 Å, respectively; Extended Data Fig. 5b). As previously described for CJF-III-288 in its complex with the gp120 core ^26^, the CJF-III-288 binding pocket includes the highly conserved S^375^ and D^368^, a continuous segment from G^472^ to R^476^, a continuous β20-β21 loop from I^424^ to I^430^, and the H^105^ of α1, which makes contacts with the propyl group of the compound (Fig. 6,A, left blow-up view). Of note, there is a high degree of similarity between the BG505 SOSIP.664 and gp120 core_e_ CJF-III-288 binding pockets (Extended Data Fig. 6).

**Fig. 6.**
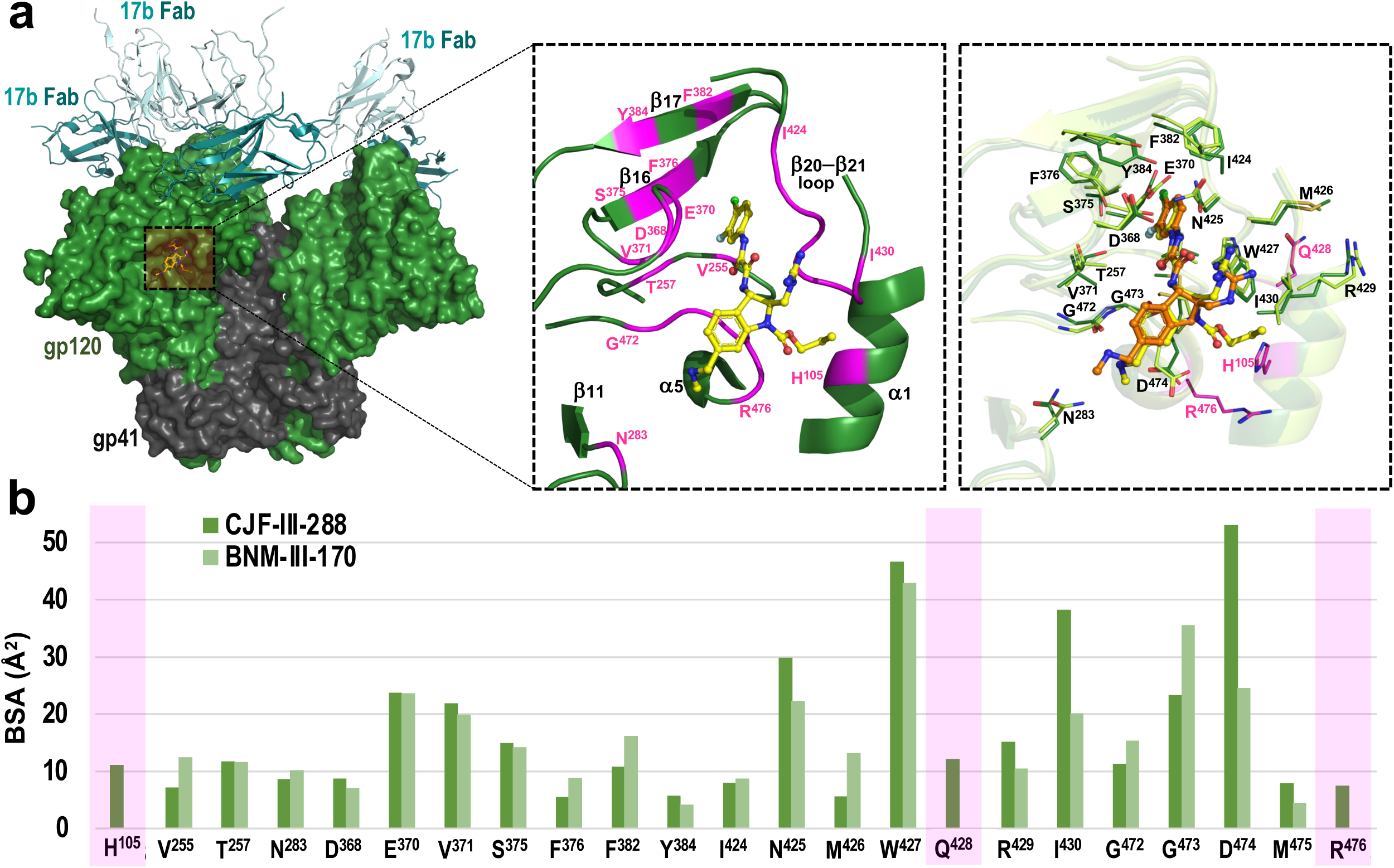
Cryo-EM structure of the CJF-III-288-BG505 SOSIP.664-17b Fab complex and a comparison of the binding pockets of CJF-III-288 and BNM-III-170 in BG505 SOSIP.664. **a**, Left, the overall structure of CJF-III-288-BG505-17b Fab complex with a molecular surface displayed over gp120 and gp41, gp120 is colored dark green and gp41 is colored grey. The 17b Fab variable region is shown as a cartoon with darker and lighter shades of cyan for heavy and light chains respectively. A side view of the complex is shown. Middle, a blow-up view shows the details of the CJF-III-288 binding pocket. Secondary structures are shown within the pocket. Residues forming the pocket are colored magenta and are labeled. Right, comparison of the two binding pockets of CJF-III-288 and BNM-III-170 bound to BG505 SOSIP.664 (as in PDB 7LO6)^44^. Pockets in protomer A in each complex were selected to do the comparison. CJF-III-288 is shown as yellow sticks and BNM-III-170 is shown as orange sticks. The gp120 of CJF-III-288 and BNM-III-170 BG505 SOSIP.664 complexes are colored darker and lighter shades of green respectively. **b,** The residue-resolved buried-surface-area (BSA) of gp120 residues contributing to the CJF-III-288-protein and BNM-III-170-protein interfaces, as determined by PISA. BSA values represent the average of the three copies in the trimer. Residues present only in the CJF-III-288-BG505 SOSIP.664 complex are highlighted with magenta.

We next compared the binding modes of CJF-III-288 and BNM-III-170 within BG505 SOSIP.664 (Fig. 6a, right blow-up view) within a single gp120 protomer. The chemical structures of BNM-III-170 and CJF-III-288 are nearly identical except for the replacement of the indane in BNM-III-170 with an indoline in CJF-III-288 and the addition of a propyl carbamate in CJF-III-288; the replacement of the indane C3 with the indoline N3 affords the addition of this distinguishing propyl carbamate. The BG505 SOSIP.664-bound conformation of CJF-III-288 and BNM-III-170 is highly similar for the identical portions of both compounds with the exception of the positions of the dimethylamine attached to the C5 carbon and the guanidinium group attached to the C2 carbon that show different orientations. As a consequence of this conformational change, I^430^ and D^474^ of gp120 contribute more buried-surface area (BSA) to the CJF-III-288-BG505 SOSIP.664-17b Fab complex interface as compared to BNM-III-170 (Fig. 6b). As expected, CJF-III-288, which is modified with a propyl-carbamat attached to the N3 nitrogen of the indoline ring, contacts three more gp120 residues in its binding interface, specifically H^105^, Q^428^ and R^476^, that are not utilized by BNM-III-170. These contacts contribute an additional 30.71 Å^2^ of BSA to the CJF-III-288 binding interface. Overall, the complexes of CJF-III-288 and BNM-III-170 with the BG505 SOSIP.664-17b Fab are formed with a BSA of 980.0 Å^2^ and 818.4 Å^2^, respectively.

Next, we examined the details of the conformation of the trimer in the CJF-III-288-BG505 SOSIP.664-17b and BNM-III-170-BG505 SOSIP.664-17b complexes (Fig. 7). When the two complexes are aligned based upon gp120/gp41 protomers, protomer B and protomer C superimpose relatively well. However, protomer A did not align well, which confirms a noticeable difference in the trimer assembly and position of the 17b Fab. In the CJF-III-288-BG505 SOSIP.664-17b Fab complex, gp120 subunits rotate away from the central gp41 helices as compared to trimer complex with BNM-III-170 (Fig. 7a,b). This rotation is evident for all three protomers to a variable extent, with the greatest rotation in protomer A. A hallmark of the trimer assembly changes includes the outward displacement of gp120 relative to the central gp41 helices which can be easily measured by the change of position of the gp120 α0 helix that packs against the gp41 HR1 helix. In the CJF-III-288 structure, the α0 helix rotates approximately 7.8 Å, 1.5 Å and 1.3 Å (for protomers A, B and C, respectively) away from the central gp41 helices as compared to the BNM-III-170 structure (Fig. 7b). This confirms the dissimilarity of the overall trimer assembly and supports the notion that the CJF-III-288-BG505 SOSIP.66417b Fab trimer is more asymmetric and more ‘open’ as compared to its BNM-III-170-triggered counterpart. Most importantly, the differences in Env trimer assembly directly define the way in which the 17b IgG approaches the Env trimer (Fig. 7 a,b). We calculated a relative angle of approach for the 17b Fab in each protomer in both structures based upon the center of mass of the 17b Fab variable domains relative to the superimposed gp41 centers (gp41 residues 570-595 of both trimers). We found that the average Fab variable chain position was different for each protomer of the trimer, again with the largest differences for protomer A (the shifts between the relative positions of 17b Fab in protomers A, B and C are 13.4 Å, 3.2 Å and 1.9 Å, respectively (Fig. 7a). 17b bound orientations are the most altered in protomer A, and include a significant displacement of the relative variable heavy and light chain orientations (Fig. 7b). Finally, we also compared the changes to the BG505 SOSIP trimer assembly induced by CJF-III-288 to those induced by other agents that target the CD4 binding site, including CD4 and other CD4mcs (Fig. 7c). The distance between the Phe43 pocket residue 375 and the angle of the gp120 protomer rotation in the CJF-III-288-BG505 SOSIP. 664-17b Fab complex differs from that in the BNM-III-170-BG505 SOSIP.664-17b complex. However, it closely resembles the asymmetric Env Conformation B of the CD4-BG505 SOSIP.664/E51 Fab complex, formed by E51, a tyrosine-sulfated CoRBS antibody that binds gp120, mimicking CCR5 interactions ^45^. This analysis demonstrates that CJF-III-288 stabilizes the Env trimer in a more asymmetric and open conformation compared to BNM-III-170, altering the angles by which 17b approaches the CoRBS in each protomer. The changes are the most pronounced in one protomer.

**Fig. 7.**
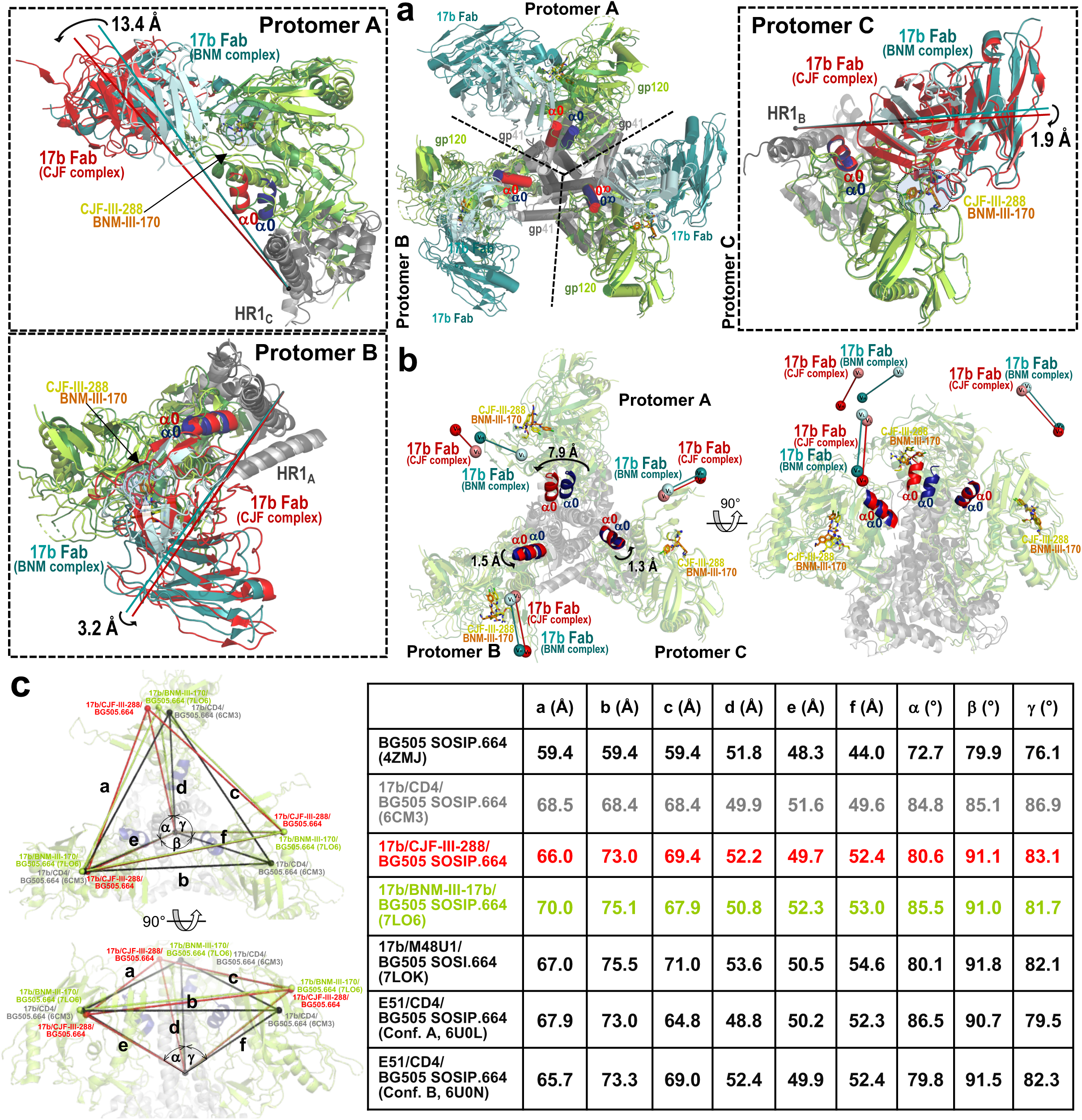
The CJF-III-288 versus the BNM-III-170-bound conformation of Env. **a**, CJF-III-288-BG505 SOSIP.664-17b Fab and BNM-III-170-BG505 SOSIP.664-17b Fab (PDB:7LO6) complexes superimposed based upon gp120/gp41 protomers. *Center*, the gp120 and gp41 in the CJF-III-288 and BNM-III-170 complexes are shown as ribbons in darker and lighter shades of green/grey, respectively (the α0 helix is shown as red/blue). *Right and left,* blow-up views show the structural alignments of Protomers A, B and C; the 17b Fab is colored light and dark cyan for the BNM-III-170 complex and red for the CJF-III-288 complex. The angle of approach for 17b in each complex is shown by a line drawn from the center of gp41 (calculated as the average of gp41 α-carbon positions for residues 570-595 in both structures) and the center of mass of the Fab variable domain (calculated as the average α-carbon position for heavy and light chain variable residues) with the distance between the Fab centers shown above by a black arrow. **b**, Changes to the opening of the timer induced by CJF-III-288 or BNM-III-170. The CJF-III-288-BG505 SOSIP.664 and BNM-III-170-BG505 SOSIP.664 trimers are shown superimposed based upon gp120/gp41 protomers (colored as in panel **a**). Relative bound positions of 17b Fab from each complex are shown with heavy-and light-chain (V_H_ and V_L_) positions determined by the center of mass of their α-carbon atoms (displayed as balls). The distance between the centers of the α0-helices in the two structures (calculated as the average of α-carbon atoms for residues 65-73) in the superimposed trimers is shown above with a black arrow indicating the rotation of the helix. **c,** Changes to overall Env trimer assembly, calculated based upon changes to the position of the α-carbon of S375 relative to the gp41 center (calculated as the average of the α-carbon positions of the central gp41 α7 helices, residues 570-595). Distances between the 375 Cα of each protomer (a, b c), the 375Cα and the gp41 center (d, e, f) and the angle between the gp41 center and two neighboring 375 Cα’s (α, β, γ) are shown.

### CryoET structure of HIV-1 Env in complex with CJF-III-288 and 17b Abs on virions confirms the asymmetric opening of Env in presence of CJF-III-288

Next, we employed CryoET to assess geometry of17b binding to Env on intact virions in the presence of CJF-III-288. AT-2 inactivated HIV-1_BaL_ virus^41,46,47^ was incubated with 100 µM CJF-III-288, and subsequently incubated on ice with 100 µg/ml 17b Ab (full IgG) before being plunge frozen and analyzed by and CryoET. 17b Ab binding to Env trimers was apparent in tomographic slices (Fig. 8a). Subtomogram averaging and classification was performed to solve the structure of Env bound to three 17b Abs (21.5 Å resolution, C1 symmetry) (Fig. 8b,c, Extended Data Fig. 7 a-f). Due to high structural heterogeneity from using full IgG, thorough classification was needed to isolate Env trimers bound to three 17b Abs (Extended Data Fig. 7 b,d). All three outwardly protruding V1/V2 loops characteristic of an open CD4-bound Env were clearly present in the average structure despite the absence of CD4 (Fig. 8b,c) ^20,41,48,49^.

**Fig. 8.**
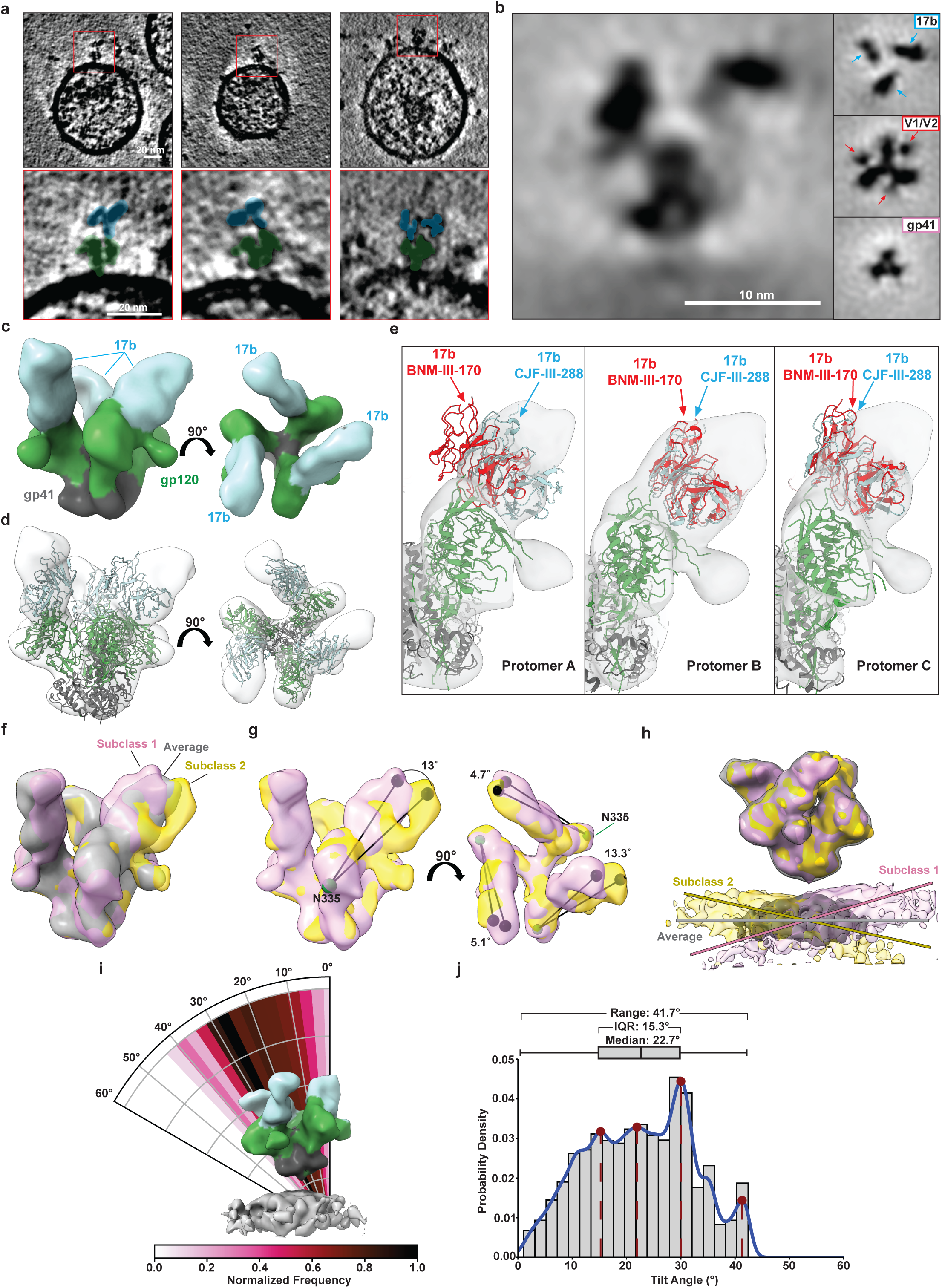
CryoET structure of HIV-1 Env in complex with CJF-III-288 and 17b Abs on virions. **a**, AT-2 inactivated HIV-1_BaL_ was incubated with 100 µM CJF-III-288 before addition of 100 µg/ml 17b Abs and subsequent plunge freezing for cryoET analysis. Tomographic slices of 17b Abs binding to Env on virions in the presence of CJF-III-288 are shown (blue: 17b, green: Env). **b**, Subtomogram averaging was performed on Env in complex with CJF-III-288 and 17b Abs (EMDB: 46646). A central slice along the density is shown. Panels to the right show three slices along the length of the structure. 17b Fabs, V1/V2 loops, and gp41 domains are indicated by arrows. **c**, Isosurface rendering of the structure. **d**, The atomic model of BG505 SOSIP.664 in complex with CJF-III-288 and 17b Fab was rigid fit into the cryoET density. **e**, The atomic model of BG505 SOSIP.664 in complex with BNM-III-170 and 17b Fab (PDB: 7LO6) ^44^ was superimposed onto the gp41 helices of the CJF-III-288 atomic model (PDB:9CF5, this paper). The three 17b Fab domains from the BNM-III-170 structure (red) and the CJF-III-288 structure (blue) are compared for their agreement with the cryoET density. **f**, Focused classification on the 17b Fab revealed heterogeneity in the 17b binding angle. Two representative subclasses are shown (pink and yellow) and compared to the combined average (grey). **g**, The 17b Fab angle between subclass 1 and subclass 2 relative to the approximate position of N^335^ (green sphere) is shown for each protomer. **h**, Focused classification on the membrane region revealed subclasses at different tilting angles. Two representative subclasses (pink and yellow) are compared with the combined average density (grey). **i**, Per-particle tilting analysis. The polar plot coloring indicates the normalized frequency of angles (n = 3512). **j**, Histogram and box-and-whisker plot showing the distribution of Env tilting. The median, range, and interquartile ranges are shown. The most prominent four apparent modes are indicated by red circles and dashed lines.

Structures of BG505 SOSIP.664 trimers bound to 17b Fab and either CJF-III-288 (PDB: 9CF5, Fig. 6 this manuscript) or BNM-III-170 CD4mc^44^ (PDB:7LO6) were compared with the cryoET density map (Fig. 8d,e). The CJF-III-288 structure was first rigidly fit into the cryoET density map before superimposing the BNM-III-170 structure by the gp41 residues. Upon examining all three 17b Fabs, the BG505 SOSIP.664 structure bound with CJF-III-288 best agreed with the cryoET density (Fig. 8e). Consistent with the analysis of differing asymmetry between the two SOSIP structures, one 17b Fab in the BNM-III-170 adopted a different orientation than the respective cryoET density (Fig. 8e).

We assessed the heterogeneity of the cryoET structure bound to three 17b molecules through focused classification. Classification on the 17b Fab with weaker density revealed subclasses with different binding orientations of 17b (Fig. 8f, Extended Data Fig. 7g). This suggests that 17b Fabs may adopt a range of conformations when bound to Env in complex with CJF-III-288 (Extended Data Movie 1), which is consistent with the relatively lower resolution observed on one 17b Fab bound to the CJF-III-288 SOSIP.664 structure (Extended Data Fig. 4) as well as the smFRET data indicating dynamics (Fig. 5). The orientation of the Fabs was found to vary by approximately 13° relative to N^355^ which is near loop V4, while the orientation of the other two Fabs remained within ∼5° of each other (Fig. 8g). Classification on the membrane region resulted in subclasses showing Env at different tilting angles (Fig. 8h and Extended Data Fig. 7h), similar to previous observations of unliganded Env tilting^50^. Subclasses and assessment of per-particle tilting distributions revealed a median Env tilt at approximately 23° on the membrane with a range of approximately 0-42° and notable modes near 15°, 22°, 30°, and 41° (Fig. 8i,j). This analysis reveals the flexibility of the Env MPER region and Fab binding orientation when Env is in complex with CJF-III-288 and 17b Ab.

## DISCUSSION

HIV-1 Env conformation significantly impacts the susceptibility of HIV-1-infected cells to ADCC. The unliganded Env trimer presents at the surface of HIV-1-infected cells predominantly assumes a “closed” State-1 conformation that protect infected cells from ADCC-mediated by nnAbs typically elicited in PLWH^16,22,23^. CD4 downmodulation by accessory proteins Nef and Vpu, along with multiple intermolecular interactions within the Env trimer helps to maintain this “closed” conformation, thus preventing the exposure of CD4i Env epitopes^18,22,23,51–53^. The use of CD4mcs to “open-up” the Env trimer and expose these vulnerable epitopes has been proposed as a new strategy to eliminate HIV-1-infected cells^25–27,31,32,37,54^. Rational, structure-based design has led to the generation of indoline CD4mcs showing broader and more potent viral inhibition and ADCC activity compared with previous indane CD4mcs^26^. Here, we report that the lead indoline CD4mc, CJF-III-288, has gained the capacity to sensitize HIV-1-infected cells to ADCC mediated by anti-CoRBS Abs, contributing to its improved ADCC activity.

This property is notable because the well-characterized anti-CoRBS Abs are unable to mediate potent ADCC against infected cells exposing either “closed” or “open” Env conformations. When combined with indane CD4mcs, anti-CoRBS Abs were reported to contribute to the ADCC response by enabling anti-cluster A Abs engage the Env trimer, at which point the Fc-portion of both families of Abs contribute to the ADCC response^31–33,54^. The exceptional ability of CJF-III-288 to enable anti-CoRBS to mediate ADCC against cells infected with multiple primary viruses, even at low concentrations of CD4mc (Figs. 1-3), underscores the therapeutic potential of the indoline CD4mcs.

The improved ADCC activity of CJF-III-288 compared with that of the indane CD4mcs likely relates to its superior ability to shift the Env conformational landscape from State 1 to open downstream conformations, which are more susceptible to ADCC mediated by nnAbs and PLWH plasma^31,53^. Compared to BNM-III-170, treatment with CJF-III-288 led to a more pronounced reduction of State 1 occupancy, even in the absence of 17b (Fig. 5). The combination of CJF-III-288 and 17b further destabilized State 1 and stabilized State 3, again to a greater extent than BNM-III-170 and 17b. The maintained conformational dynamics of Env bound to CJF-III-288 and 17b likely contributes to the variability in 17b orientation seen in our structural analyses. Although CJF-III-288 slightly increased the occupancy of State 2A, under no condition did State 2A become predominant as previously seen in the presence of anti-cluster A Abs^31^. This indicates that State 3 is also a viable substrate for ADCC. The asymmetry in Env conformation revealed through our structural analyses are likely of insufficient amplitude to detect by smFRET with the current fluorophore attachments sites. Overall, our findings suggest that CJF-III-288’s ability to stabilize State 3 contributes to its activity in sensitizing HIV-1-infected cells to ADCC mediated by anti-CoRBS and PLWH plasma.

Structural analyses helped delineate the mechanism behind the improved ADCC activity of anti-CoRBS Abs in the presence of CJF-III-288. The structure of the CJF-III-288-BG505 SOSIP.664-17b complex obtained by cryo-EM revealed significant differences in the trimer assembly relative to the previously published BNM-III-17-BG505 SOSIP.664-17b complex^44^ (Fig. 7). This is evidenced by a substantial outward rotation of the gp120 subunits, particularly in protomer A, where the α0 helix shifts significantly further away from the central gp41 helices compared to the BNM-III-170 complex. The CJF-III-288-BG505 SOSIP.664-17b closely resembles the asymmetric Env conformation B of the CD4-BG505 SOSIP.664/E51 Fab complex^45^. E51 is a tyrosine-sulfated CoRBS antibody that was shown to induce an asymmetric opening of the Env trimer leading to rearrangements of relative positions of gp120/gp41 protomers within the Env trimer. It has been suggested that E51-bound Env conformations represent structural intermediates of Env, wherein the CoRBS is being formed before transitioning into a fully open state. Thus, our data suggest that CJF-III-288 is able, on its own, to promote the conformational changes required to form the coreceptor binding site. Indeed, both indane and indoline CD4mcs must be able to induce coreceptor binding, based on activation of HIV-1 infection of CD4-negative, coreceptor-expressing cells ^26,55^. However, BNM-III-170 and CJF-III-288 induce distinct asymmetric conformations in the Env trimer, orienting the CoRBS differently. These structural distinctions would be expected to influence how the anti-CoRBS Abs approach one of the gp120 subunits of the trimer (Fig. 7). Although changes in the Ab approach were noticed for each gp120 protomer in the trimer, the largest deviation was observed for one (protomer A) where a shift of 13.4 Å for the center of mass of the Fab variable domain of the anti-CoRBS Ab was detected. Cryo-ET analysis of the membrane-bound Env trimer confirmed the asymmetric opening of the trimer, as the CJF-III-288-BG505 SOSIP.664-17b structure fits relatively well into the cryo-ET density map (Fig. 8). Assessment of the heterogeneity of the cryo-ET structure revealed the flexibility of the Env MPER and anti-CoRBS Ab binding when Env is in complex with CJF-III-288. CJF-III-288 may allow anti-CoRBS Fabs to adopt a range of conformations, notably with a variation of 13° within the protomer A, consistent with cryo-EM findings. Furthermore, the observed variation of the Env trimer’s tilting angle also has the potential to facilitate Fc receptor engagement on effector cells by accommodating diverse antibody/Env orientations, thereby improving alignment with the Fc receptor’s angle of approach.

Overall, both functional and structural data strongly support the notion that CJF-III-288 induces open, asymmetric Env conformations that enable anti-CoRBS Abs to bind in orientations that allow for Fc receptor engagement and ADCC. This highlights CJF-III-288’s potential as a therapeutic agent, capable of harnessing the full potential of non-neutralizing CD4i antibody responses to target HIV-1-infected cells through Fc-effector activities.

## ONLINE METHODS

### Ethics Statement

Written informed consent was obtained from all study participants and research adhered to the ethical guidelines of CRCHUM and was reviewed and approved by the CRCHUM institutional review board (ethics committee, approval number MP-02-2024-11734). Research adhered to the standards indicated by the Declaration of Helsinki.

### Cell lines and primary cells

293T human embryonic kidney cells (obtained from ATCC) were maintained at 37°C under 5% CO2 in Dulbecco’s Modified Eagle Medium (DMEM) (Wisent, St. Bruno, QC, Canada), supplemented with 5% fetal bovine serum (FBS) (VWR, Radnor, PA, USA) and 100 U/mL penicillin/streptomycin (Wisent). Human peripheral blood mononuclear cells (PBMCs) from 5 HIV-negative individuals (4 males, 1 female) and 4 PLWH (4 males) obtained by leukapheresis and Ficoll-Paque density gradient isolation were cryopreserved in liquid nitrogen until further use. CD4+ T lymphocytes were purified from resting PBMCs by negative selection using immunomagnetic beads per the manufacturer’s instructions (StemCell Technologies, Vancouver, BC) and were activated with phytohemagglutinin-L (10 µg/mL) for 48 h and then maintained in RPMI 1640 (Thermo Fisher Scientific, Waltham, MA, USA) complete medium supplemented with rIL-2 (100 U/mL).

### Antibody production

FreeStyle 293F cells (Thermo Fisher Scientific) were grown in FreeStyle 293F medium (Thermo Fisher Scientific) to a density of 1 × 10^6^ cells/mL at 37°C with 8% CO2 with regular agitation (150 rpm). Cells were transfected with plasmids expressing the light and heavy chains of each mAb using ExpiFectamine 293 transfection reagent, as directed by the manufacturer (Thermo Fisher Scientific). One week later, the cells were pelleted and discarded. The supernatants were filtered (0.22-μm-pore-size filter), and antibodies were purified by protein A affinity columns, as directed by the manufacturer (Cytiva, Marlborough, MA, USA). Antibodies were dialyzed against phosphate-buffered saline (PBS) and stored in aliquots at −80°C. To assess purity, antibodies were loaded on SDS-PAGE polyacrylamide gels in the presence or absence of β-mercaptoethanol and stained with Coomassie blue. The anti-CoRBS 17b Ab Fab fragments were prepared from purified IgG (10 mg/mL) by proteolytic digestion with immobilized papain (Pierce, Rockford, IL) and purified using protein A, followed by gel filtration chromatography on a Superdex 200 16/60 column (Cytiva).

### Plasmids and proviral constructs

Transmitted/founder infectious molecular clones (IMCs) of patient CH058 was inferred, constructed, and biologically characterized as previously described^56,57^. IMCs encoding HIV-1 reference strains JR-FL, JR-CSF, AD8 (Vpu+) and BG505 (T332N) were described elsewhere ^58–61^. The vesicular stomatitis virus G (VSV-G)-encoding plasmid was previously described ^62^.

### Viral production, infections and *ex vivo* amplification

For *in vitro* infection, vesicular stomatitis virus G (VSV-G)-pseudotyped HIV-1 viruses were produced by co-transfection of 293T cells with an HIV-1 proviral construct and a VSV-G-encoding vector using the calcium phosphate method. Two days post-transfection, cell supernatants were harvested, clarified by low-speed centrifugation (300 × g for 5 min), and concentrated by ultracentrifugation at 4°C (100,605 × g for 1 h) over a 20% sucrose cushion. Pellets were resuspended in fresh RPMI, and aliquots were stored at −80°C until use. Viral preparations were titrated directly on primary CD4+ T cells to achieve similar levels of infection among the different IMCs tested. Viruses were then used to infect activated primary CD4+ T cells from HIV-1 negative donors by spin infection at 800 × *g* for 1 h in 96-well plates at 25 °C. All experiments using VSV-G-pseudotyped HIV-1 isolates were done in a biosafety level 3 laboratory following manipulation protocols accepted by the CRCHUM Biosafety Committee, which respects the requirements of the Public Health Agency of Canada.

To expand endogenously infected CD4+ T cells, primary CD4+ T cells were isolated from PBMCs obtained from PLWH by negative selection as described above. Purified CD4+ T cells were activated with PHA-L at 10 μg/mL for 48 h and then cultured for at least 6 days in RPMI 1640 complete medium supplemented with rIL-2 (100 U/ml) to reach greater than 5% infection for the ADCC assay.

### Antibodies and human plasma

The following anti-Env Abs were used as primary mAbs to stain HIV-1-infected primary CD4+ T cells: anti-cluster A A32, N5i5; anti-co-receptor binding site 17b, C2, N12i2, 48D, 412D, X5; anti-gp120 outer domain 2G12. Goat anti-human IgG (H+L) (Thermo Fisher Scientific) pre-coupled to Alexa Fluor 647 were used as secondary antibodies in flow cytometry experiments. The A32 mAbs was conjugated with Alexa-Fluor 647 (Thermo Fisher Scientific) as per the manufacturer’s protocol and was also used for cell-surface staining of HIV-1-infected cells. Mouse anti-human CD4 (Clone OKT4, FITC-conjugated; Biolegend, San Diego, CA, USA) and anti-p24 mAb (clone KC57; PE-Conjugated; Beckman Coulter) or Mouse anti-human CD4 (Clone OKT4, PE-conjugated; Biolegend, San Diego, CA, USA) and anti-p24 mAb (clone KC57; FITC-Conjugated; Beckman Coulter) were used to identify the productively-infected cells as previously described^24^. Plasma from PLWH, obtained from the FRQS-AIDS and Infectious Diseases Network (the Montreal Primary HIV Infection Cohort ^63,64^), were collected, heat-inactivated, and conserved as previously described^27^.

### Small CD4-mimetics

The small-molecule CD4-mimetic compounds (CD4mc) BNM-III-170, CJF-III-288 and TFH-I-116-D1 were synthesized as described previously^26,37^. The compounds were dissolved in dimethyl sulfoxide (DMSO) at a stock concentration of 10 mM and diluted in phosphate-buffered saline (PBS) for cell-surface staining or in RPMI-1640 complete medium for ADCC assays.

### Flow cytometry analysis of cell-surface staining

Cell surface staining was performed at 48h post-infection. Mock-infected or HIV-1-infected primary CD4+ T cells were incubated for 45 min at 37°C with anti-Env mAbs (5 µg/mL) or plasma from PLWH (1:1000) in the presence or absence of different concentrations of CD4mc (BNM-III-170, TFH-I-116-D1, CJF-III-288) or equivalent volume of DMSO. Cells were then washed twice with PBS and stained with the appropriate Alexa Fluor 647-conjugated secondary antibody (2 µg/mL) and the viability dye staining (Aqua Vivid; Thermo Fisher Scientific) for 20 min at room temperature. Cells were then stained with FITC-or PE-conjugated mouse anti-CD4 Abs. After two PBS washes, cells were fixed in a 2% PBS-formaldehyde solution. Infected cells were then permeabilized using the Cytofix/Cytoperm Fixation/ Permeabilization Kit (BD Biosciences, Mississauga, ON, Canada) and stained intracellularly using PE or FITC-conjugated mouse anti-p24 mAb ( 1:100 dilution). The percentage of productively-infected cells was determined by gating on the living CD4^low^p24^high^ cell population as previously described ^24^. Samples were acquired on an FORTESSA flow cytometer (BD Biosciences), and data analysis was performed using FlowJo v10.5.3 (Tree Star, Ashland, OR, USA). Area under the curve (AUC) values were calculated from the median fluorescence intensities obtained at different CD4mc concentrations (log10-transformed) using GraphPad Prism software.

### Antibody-dependent cellular cytotoxicity (ADCC) assay

Measurement of ADCC using a fluorescence-activated cell sorting (FACS)-based infected cell elimination (ICE) assay was performed at 48 h post-infection. Briefly, HIV-1-infected primary CD4+ T cells were stained with AquaVivid viability dye and cell proliferation dye eFluor670 (Thermo Fisher Scientific) and used as target cells. Cryopreserved autologous PBMC effectors cells, stained with cell proliferation dye eFluor450 (Thermo Fisher Scientific), were added at an effector: target ratio of 10:1 in 96-well V-bottom plates (Corning, Corning, NY). Target cells were treated with CD4mc (CJF-III-288, BNM-III-170 or TFH-I-116-D1) at indicated concentrations or equivalent volume of DMSO. Anti-Env mAbs (5 µg/mL) or PLWH plasma (1:1000 dilution) were added to appropriate wells and cells were incubated for 5 min at room temperature. For blocking experiments, infected cells were pre-incubated with the anti-CoRBS Ab 17b Fab fragment at 10µg/mL in the presence of CD4mc before adding PLWH plasma. The plates were subsequently centrifuged for 1 min at 300 × g, and incubated at 37 °C, 5 % CO_2_ for 5 h. before being stained with FITC-or PE-conjugated Mouse anti-CD4 Abs. After one PBS wash, cells were fixed in a 2% PBS-formaldehyde solution. Infected cells were then permeabilized using the Cytofix/Cytoperm Fixation/ Permeabilization Kit (BD Biosciences, Mississauga, ON, Canada) and stained intracellularly using PE or FITC-conjugated mouse anti-p24 mAb (1:100 dilution). Productively-infected cells were identified based on p24 and CD4 detection as described above. Samples were acquired on an FORTESSA flow cytometer (BD Biosciences), and data analysis was performed using FlowJo v10.5.3 (Tree Star, Ashland, OR, USA). The percentage of ADCC was calculated with the following formula: [(% of CD4^low^p24^+^ cells in Targets plus Effectors) − (% of CD4^low^p24^+^ cells in Targets plus Effectors plus plasma or mAbs) / (% of CD4^low^p24^+^ cells in Targets) × 100] by gating on infected lived target cells. Area under the curve (AUC) values were calculated from the percentage of ADCC obtained at different CD4mc concentrations (log10-transformed) using GraphPad Prism software.

### smFRET imaging

For smFRET imaging, pseudovirions were formed with the HIV-1_NL4-3_ core and HIV-1_JR-FL_ Env. To facilitate fluorophore attachment, an amber stop codon was introduced at position N135 in the V1 loop by site-directed mutagenesis, and the A4 peptide was inserted into the V4 loop of Env by overlap-extension PCR, as described ^29^. Briefly, HEK-293T FIRB cells^65^ were co-transfected with plasmids encoding an aminoacyl tRNA synthetase and corresponding suppressor tRNA (NESPylRS^AF^/hU6tRNA^Pyl^) ^66^, eRF1-E55D, pNL4-3 ΔRT ΔEnv ^16^, and a 20:1 mass ratio of wild-type HIV-1_JR-FL_ Env to tagged Env. Pseudovirus was collected 48 hours post-transfection and pelleted through a 10% sucrose cushion in PBS by ultracentrifugation for 2 h at 35,000 RPM. Pellets was then resuspended in labeling buffer (50 mM HEPES pH 7.0, 10 mM CaCl2, 10 mM MgCl2) and incubated overnight at room temperature with 5 μM LD650-coenzyme A (Lumidyne Technologies, New York,NY, USA), and 5 μM acyl carrier protein synthase (AcpS), which attaches the fluorophore to the A4 peptide in V4 of gp120, as described ^16,31,42^. The virus was then incubated with 0.5 µM Cy3-tetrazine (Jena Biosciences, Jena, Germany) for 30 min at room temperature, followed by addition of 60 μM DSPE-PEG2000-biotin (Avanti Polar Lipids, Alabaster, AL, USA) and incubation for an additional 30 min at room temperature. Finally, virus was purified by ultracentrifugation on a 6–30% OptiPrep (Sigma-Aldrich, MilliporeSigma, Burlington, MA, USA) density gradient for 1 hour at 35,000 RPM at 4 °C using a SW40Ti rotor (Beckman Coulter Life Sciences, Brea, CA, USA). Gradient fractions were collected, analyzed by western blot, aliquoted, and stored at –80°C until their use in imaging experiments.

Where indicated, labelled pseudovirions were incubated with 50 µg/ml 17b mAb and 100 μM CD4mc for 1 hour at room temperature, followed by immobilization on streptavidin-coated quartz slides. Virions were then imaged on a custom-built wide-field prism-based TIRF microscope ^67,68^. Imaging was performed in PBS pH 7.4, containing 1 mM trolox (Sigma-Aldrich, St. Louis, MO, USA), 1 mM cyclooctatetraene (COT; Sigma-Aldrich, St. Louis, MO, USA), 1 mM 4-nitrobenzyl alcohol (NBA; Sigma-Aldrich, St. Louis, MO, USA), 2 mM protocatechuic acid (PCA; Sigma-Aldrich, St. Louis, MO, USA), and 8 nM protocatechuate 3,4-deoxygenase (PCD; Sigma-Aldrich, St. Louis, MO, USA) to stabilize fluorescence and remove molecular oxygen. Where indicated, concentrations of mAb 17b and CD4mc were maintained during imaging. smFRET data were collected using Micromanager v2.0 ^69^at 25 frames/sec, processed, and analyzed using the SPARTAN software (https://www.scottcblanchardlab.com/software) in Matlab (Mathworks, Natick, MA, USA) ^70^. smFRET traces were identified according to criteria previously described^31^, and traces meeting those criteria were verified manually. FRET histograms were generated by compiling traces from each of three technical replicates and the mean probability per histogram bin ± standard error was calculated. Traces were idealized to a five-state HMM (four nonzero-FRET states and a zero-FRET state) using the maximum point likelihood (MPL) algorithm^71^ implemented in SPARTAN as described^31^. The idealizations were used to determine the occupancies (fraction of time until photobleaching) in each FRET state, and construct Gaussian distributions, which were overlaid on the FRET histograms to visualize the results of the HMM analysis. The FRET state occupancies for each trajectory were used to construct violin plots in Matlab, as well as calculate mean and median occupancies, standard errors, and quantiles.

### Protein expression and purification for Cryo-EM studies

BG505 SOSIP.664 was expressed in Expi293F GnTI-cells (Thermo Fisher Scientific, Catalog# A39240). One day prior to transfection, cells were diluted to a density of 1 × 10^6^ cells/ml for a total volume of 90 ml. 50 μg BG505 SOSIP.664 plasmid and 10 μg furin plasmid were mixed and diluted into 5 ml Expi293™ Expression Medium (Thermo Fisher Scientific, Catalog# A1435101). 75 μl FectoPRO (Polyplus) transfection reagent was diluted into 5 ml Expi293™ Expression Medium. Plasmids and transfection reagent were then mixed and incubated at room temperature for 10 minutes prior to addition to the cells. Cells were then grown for another 5 days, and the supernatant harvested by centrifugation. The supernatant was filtered through a 0.22 μm membrane and BG505 SOSIP.664 was purified with a PGT145 affinity column (PGT145 IgG covalently linked to Protein A agarose). Briefly, supernatant was loaded onto the PGT145 affinity column, the column then washed with PBS, and finally BG505 SOSIP.664 protein eluted with 3 M MgCl_2_. The eluted BG505 SOSIP.664 protein was immediately buffer exchanged into PBS. BG505 SOSIP.664 was further purified with a Superdex 200 10/300 GL gel filtration column (Cytiva).

17b IgG was expressed in Expi293F cells (Thermo Fisher Scientific, Catalog# A14528) with a transfection protocol similar to that of BG505 SOSIP.664. For each 100 ml volume transfection, 25 μg 17b heavy chain plasmid and 25 μg 17b light chain plasmid were mixed and diluted with 5 ml Expi293™ Expression Medium (Thermo Fisher Scientific, Catalog# A1435101). 75 μl transfection reagent FectoPRO (Polyplus) was diluted in 5 ml Expi293™ Expression Medium. Plasmids and transfection reagent were mixed and incubated at room temperature for 10 minutes before being added to 90 ml cells. Cells were grown for an additional 7 days and then supernatant was harvested by centrifugation. The supernatant was filtered through a 0.22 μm membrane and 17b IgG was purified with protein A affinity column. Briefly, the supernatant was loaded onto the protein A column, the column washed with PBS, and 17b IgG was eluted with 0.1 M glycine-HCl, pH 2.7. Fabs were generated from an overnight papain digestion of IgG at 37 °C using immobilized papain agarose (Thermo Fisher Scientific). The resulting Fab was separated from Fc and undigested IgG by passage over protein A resin. Fab was further purified by gel filtration using a Superdex 200 10/300 GL column (GE Healthcare) before use in cryo-EM sample preparation.

### Cryo-EM sample preparation

CJF-III-288 and BG505 SOSIP.664 trimer were mixed at a molar ratio of 10:1, then incubated at room temperature overnight. 17b Fab was added on the next day at a 10:1 molar ratio to BG505 SOSIP.664 and then incubated at room temperature for 4 hours. The CJF-III-288-BG505 SOSIP.664-17b complex was purified on a Superdex 200 10/300 GL column (GE Healthcare) and fractions containing CJF-III-288-BG505-17b complex were concentrated to 1.5 mg/ml. n-dodecyl-β-D-maltopyranoside (DDM) was added to a final concentration of 0.091 mM prior to freezing. To prepare cryo-specimen grids, a 3 μL aliquot of each sample was applied onto a Quantifoil 400-mesh Cu R1.2/1.3 grid with a2nm carbon fil over the holes (400-mesh Cu R1.2/1.3-2nm, Quantifoil Micro Tools GmbH), which had been freshly glow-discharged for 15 s at 15 mA using a PELCO easiGLOW (TED PELLA). The sample was then plunged into liquid ethane using a Leica GP2 (Leica Biosystems) after blotting for 5 s at 4 °C under 95% relative humidity.

### Cryo-EM data collection and processing

Data was acquired on a FEI Titan Krios electron microscope operating at 300 kV equipped with Gatan Bioquantum Image filter-K3 direct electron detector (Gatan Inc) with 20 eV energy slit. 50-frame dose fractionated movies in super resolution mode were collected at a nominal magnification of 105,000 corresponding to a calibrated physical pixel size of 0.832 Å/px (0.416 Å/px super resolution), with a total exposure dose of 54.2 e−/ Å^2^. Automated data acquisition was done in SerialEM version 4.0.27^72^. CryoSPARC ver.4.4.1^73^ was used to process the cryo-EM data. A total of 9458 movies were imported. After Patch Motion Correction and Patch CTF Estimation, 9431 micrographs were used for the following processing. An initial 150,000 particles were picked using the Blob Picker and then subjected to 2D Classification. Particles with side views were used as templates for Template Picking. An extraction box size of 400-pix was used to extract particles. After several rounds of 2D Classification, 215,095 particles with balanced side and top views were used to do Ab-initio Reconstruction. 7,078 junk particles were used to do Ab-initio Reconstruction and generate 5 junk models. The good Ab-initio model was low-pass filtered to 30 Å. Heterogeneous Refinement using the low-pass filtered good Ab-initio model and the 5 junk models as references was conducted several times to clean extracted particles. After Heterogeneous Refinement, 1,087,694 particles were used to do Non-uniform Refinement which generated a 3.5 Å density map. C1 was applied to all reconstruction and refinement steps.

### Atomic model building and refinement

An initial fit to the CryoEM reconstruction was done using the BNM-III-170-BG505 SOSIP.664-17b complex as a starting model (PDB ID 7LO6)^44^. Briefly individual protomers were fit manually to the reconstruction in ChimeraX^74^. CJF-III-288 was then added to the model by superposition of the gp120 complex of CJF-III-288 (PDB ID 8FM3) onto each protomer of the trimer. The initial model was then manually fit into the reconstruction using COOT ^75,76^ and then refined with real space refinement in PHENIX ^77,78^. Several rounds of refinement and model building were done to finalize the model. Data collection and refinement statistics are located in Extended Data Table 1.

### CryoET Sample Preparation

AT-2 inactivated HIV-1_BaL_ stocks ^46^ were incubated at room temperature for 1 hour in the presence of 100 μM CJF-III-288. The virus was transferred to ice and 100 μg/ml 17b antibody was subsequently added for an approximately 30-minute incubation. Copper Quantifoil holey grids (2/1, 200 mesh) were glow discharged for 35 seconds at 25 mA and pre-blotted with 3 μl BSA-colloidal gold solution before adding 5 μl of the virus sample preparation. Grids were blotted from the front with filter paper and plunged into liquid ethane with a homemade gravity plunger.

### CryoET Data Acquisition

CryoET grids containing plunge-frozen HIV_BaL_ with 17b IgG and CJF-III-288 were imaged on a Titan Krios electron microscope (Thermo Fisher Scientific) equipped with a 300 keV field emission gun, K3 Summit direct electron detector (Gatan), Volta Phase Plate (to enhance contrast), and a Gatan Imaging Filter. Data was collected using SerialEM software ^72^ at 64,000X magnification (1.346 Å/pixel) using a single axis bidirectional tilt series scheme from –60° to 60° with a tilt step of 3° and a cumulative specimen dose of 120 e^-^ /Å^2^. Dose fractionation mode was used to generate 9 frames per image for subsequent motion correction.

### Tomogram Reconstruction

MotionCor2 was used to process dose-fractionated images, which were then compiled into image stacks ^79^. Motion-corrected image stacks were aligned with IMOD/Etomo using gold fiducial markers ^72^. For visualization and particle picking, aligned stacks were binned by 8x or 4x and IMOD was used to generate tomograms with weighted back-projection followed by deconvolution filtering. CTFPlotter was used to estimate defocus and phase values ^80^.

### Subtomogram averaging

Subtomogram averaging was performed using PEET ^81^. HIV-1_BaL_ Env trimers bound with CJF-III-288 and 17b Abs were manually selected from tomograms binned by 8 using IMOD. A point was placed on each Env trimer and at the center of each virion. PEET program SpikeInit was then used to generate a rough estimate of the particle orientations. Particles binned by 4 were first aligned using no translational or angular searching to generate an initial formless reference of a ball sitting on a membrane surface. The membrane and Ab domains not connected to trimers (Fc domains or free Fabs domains) were subsequently masked out for further alignment. An initial model was generated with resolution limited to 45 Å by strong lowpass filtration and binning before randomly splitting the data into two halves for gold-standard refinement. PCA-based classification was used to remove junk particles and separate out two vs. three 17b-bound trimers by masking on the Fabs with noticeably weaker density ^82^. Env trimers bound to three 17b Abs were further analyzed. Small 2x-binned 3D subvolumes containing individual Env particles were reconstructed with CTF correction using IMOD program subtomosetup. One final refinement iteration was performed on these 2x binned particles. Fourier shell correlation (FSC) curves were calculated using Relion ^83^ and plotted in Python (matplotlib and scipy libraries) to estimate resolution using FSC = 0.143.

### Heterogeneity analysis of 17b Fab orientation and Env tilting

Focused PCA-based classification was performed on the final Env structure bound to three 17b Abs to assess heterogeneity using PEET ^81,82^. To determine heterogeneity in binding orientation, a spherical mask was drawn on the weakest density 17b Fab domain. Env tilting was assessed using an ellipsoid mask drawn on the membrane and MPER region. Structures were visualized in ChimeraX ^74^ and different orientations were displayed as a movie using the ChimeraX “vop morph” function.

Trimers bound to three 17b molecules were assessed for tilting on a per-particle basis. The angle between the Env orientation vector (determined through subtomogram averaging) and a vector perpendicular to the membrane at each respective Env position was determined as the Env tilting angle. To calculate the vector perpendicular to the membrane, focused angular refinement on the MPER-membrane region was performed, resulting in a disordered Env ectodomain and a well-defined membrane density. Tilt angles distributions were plotted as a polar plot in Python (ver 3.8.5, matplotlib and numpy packages) and as a histogram using R (ver. 4.3.2, ggplot2 package). The four most prominent modes were detected using the “multimode” R package (doi:10.18637/jss.v097.i09).

### Statistical analysis

Statistics were analyzed using GraphPad Prism version 9.1.0 (GraphPad, San Diego, CA, USA). Every data set was tested for statistical normality and this information was used to apply the appropriate (parametric or nonparametric) statistical test. P values <0.05 were considered significant; significance values are indicated as * P<0.05, ** P<0.01, *** P<0.001, **** P<0.0001.

## Acknowledgements

The authors thank the CRCHUM BSL3 and Flow Cytometry Platforms for technical assistance, Mario Legault from the FRQS AIDS and Infectious Diseases network for providing the human PBMCs and plasma. We thank Dennis Burton (The Scripps Research Institute) Julie Overbaugh (Fred Hutchinson Cancer Research Center) Frank Kirchhoff (Ulm University Medical Center) and Beatrice H. Hahn (University of Pennsylvania) for kindly providing the infectious molecular clone (IMC) JR-FL, BG505 (T332N), AD8 (Vpu+) and CH058TF respectively. We thank Shenping Wu (Yale CryoEM Resource Center) and John Heumann (University of Colorado Boulder) for technical and conceptual advice regarding cryoET data collection and analysis. We thank James Robinson for providing the plasmids to produce the A32 and 17b Abs. We thank NIH Intramural CryoEM Consortium (NICE) for microscopy resource and consultant. We thank John P. Moore (Cornell University) for kindly giving us BG505 SOSIP.664 plasmid and Andrew Ward (Scripps research) for providing the plasmids to produce the PGT145 Abs. This study was supported by a CIHR Team grant #422148, a Canada Foundation for Innovation (CFI) grant #41027 to A.F and by the National Institutes of Health to A.F. (R01AI148379, R01AI176531), A.F. and J.B.M. (R01AI150322), A.F. and M.P. (R01AI174908,), A.F., M.P., and A.B.S. (R01AI186809), M.P. (P01AI162242), W.M. (R37AI150560, R01AI176904, P01AI150471), and M.W.G. (F31 AI176650). Support for this work was also provided by UM1AI164562 (ERASE) to A.F., J.S. and A.B.S. A.F. was supported by a Canada Research Chair on Retroviral Entry RCHS0235 950–232424. This research was also supported by the Intramural Research Program of the *Eunice Kennedy Shriver* National Institute of Child Health and Human Development to D.M. (ZIA HD008998). G.B.B. M.B, E.B and K.D were supported by CIHR doctoral fellowships. E.B. was supported by a FRQS doctoral fellowship. M.W.G was supported by NIH T32 AI055403 and is a recipient of the Gruber Science Fellowship. The funders had no role in study design, data collection and analysis, decision to publish, or preparation of the manuscript.

## Data availability

The data that support this study are available from the corresponding authors upon request. Cryo-EM and Cryo-ET maps have been deposited in the Electron Microscopy Data Bank (EMDB) under accession codes EMD-45530 and EMD-46646, respectively. Raw movies will be uploaded to the Electron Microscopy Public Image Archive (EMPIAR). The atomic coordinates for the CJF-III-288-BG505 SOSIP.664-17b Fab complex have been deposited in the Protein Data Bank (PDB) under accession code 9CF5.

## Author contributions

Conceptualization: J.R., M.W.G, L.N, M.A.D-S, W.M., J.B.M., M.P. and A.F. Methodology: J.R., M.W.G, L.N, M.A.D-S, W.D.T., W.L., Z.C.L., A.J.M., R.K.H., D.M., W.M., J.B.M., M.P. and A.F. Investigation: J.R., M.W.G, L.N, M.A.D-S, J.B.M., W.D.T, L.M, F.Z., C.B., R.K.H., Resources: J.R., L.N, M.W.G, M.A.D-S, D.Y., L.M., T.J.C., H-C.C., M.B., G.B-B., K.D., E.B., D.C., H.M., W.A.H., J.S., A.B.S., D.M.,W.M., J.M., M.P. and A.F. Supervision: J.R., W.A.H., J.S., A.B.S., W.L., D.M., W.M., J.B.M., M.P. and A.F. Funding acquisition: W.A.H., J.S., A.B.S., J.B.M., W.M., M.P. and A.F. Writing original draft: J.R., M.W.G, L.N, M.A.D-S, W.M., J.B.M., M.P. and A.F. Writing & editing: review: J.R., L.N, M.W.G, M.A.D-S, W.D.T., L.M, F.Z., C.B., D.Y., T.J.C., H-C.C., M.B., G.B-B.,S.G., W.L., K.D., E.B., D.C., H.M., W.A.H., J.S., R.K.H., D.M., A.B.S., W.M., J.B.M., M.P. and A.F.

## DISCLAIMER

The views expressed in this manuscript are those of the authors and do not reflect the official policy or position of the Uniformed Services University, US Army, the Department of Defense, or the US Government.

## FIGURE LEGENDS

**Extended data Fig. 1.**
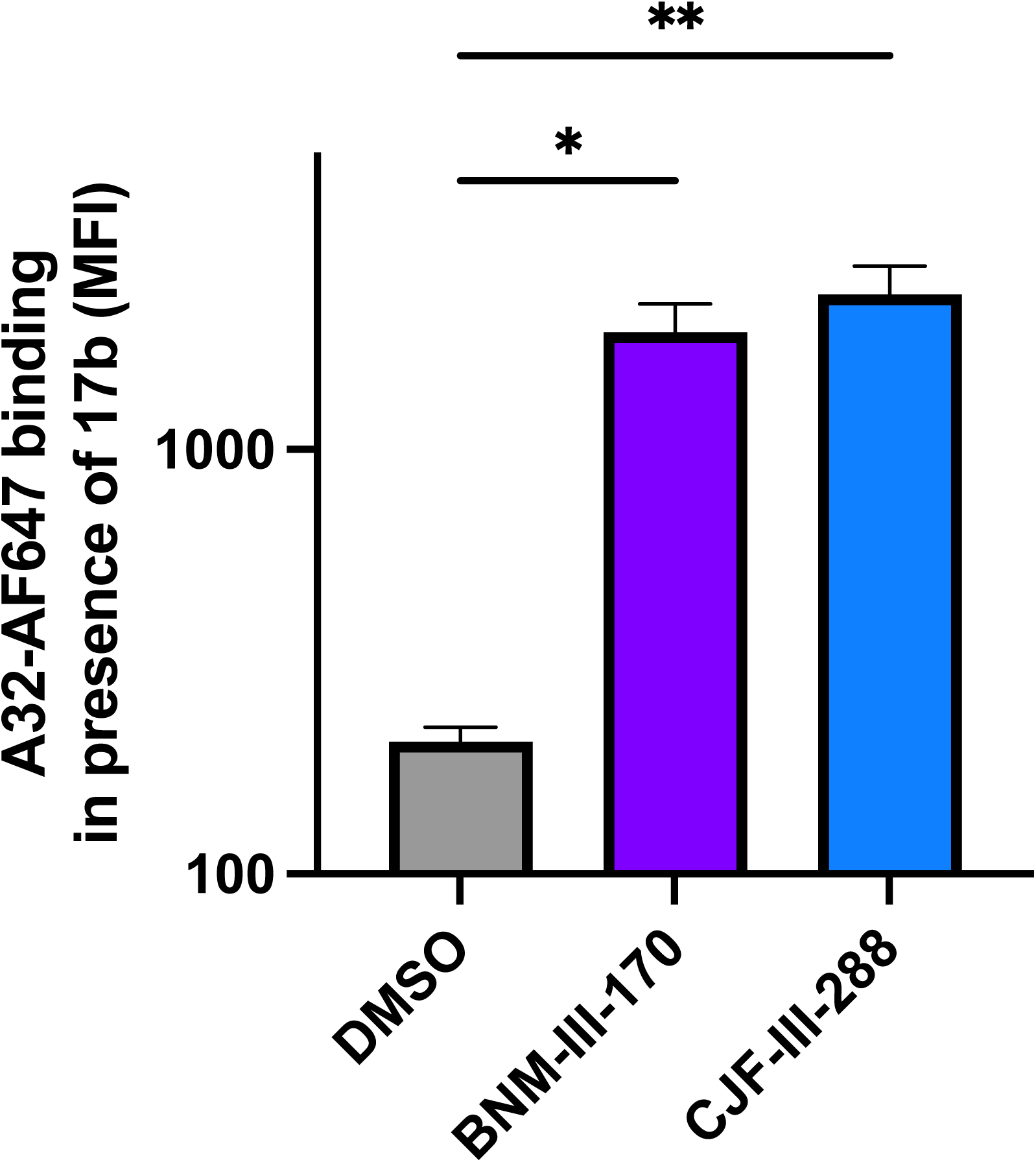
CJF-III-288 and 17b combination enable recognition of HIV-1-infected cells by A32. Recognition of HIV-1_CH058TF_-infected primary CD4+ T cells by Alexa-Fluor-647 conjugated A32 in combination with 17b and DMSO, BNM-III-170 or CJF-III-288. Graphs represent the median fluorescence intensities (MFI) obtained in 3 independent experiments. Statistical significance was tested using One-way ANOVA test with a Tukey post-test (* p<0.05,** p<0.01).

**Extended data Fig. 2.**
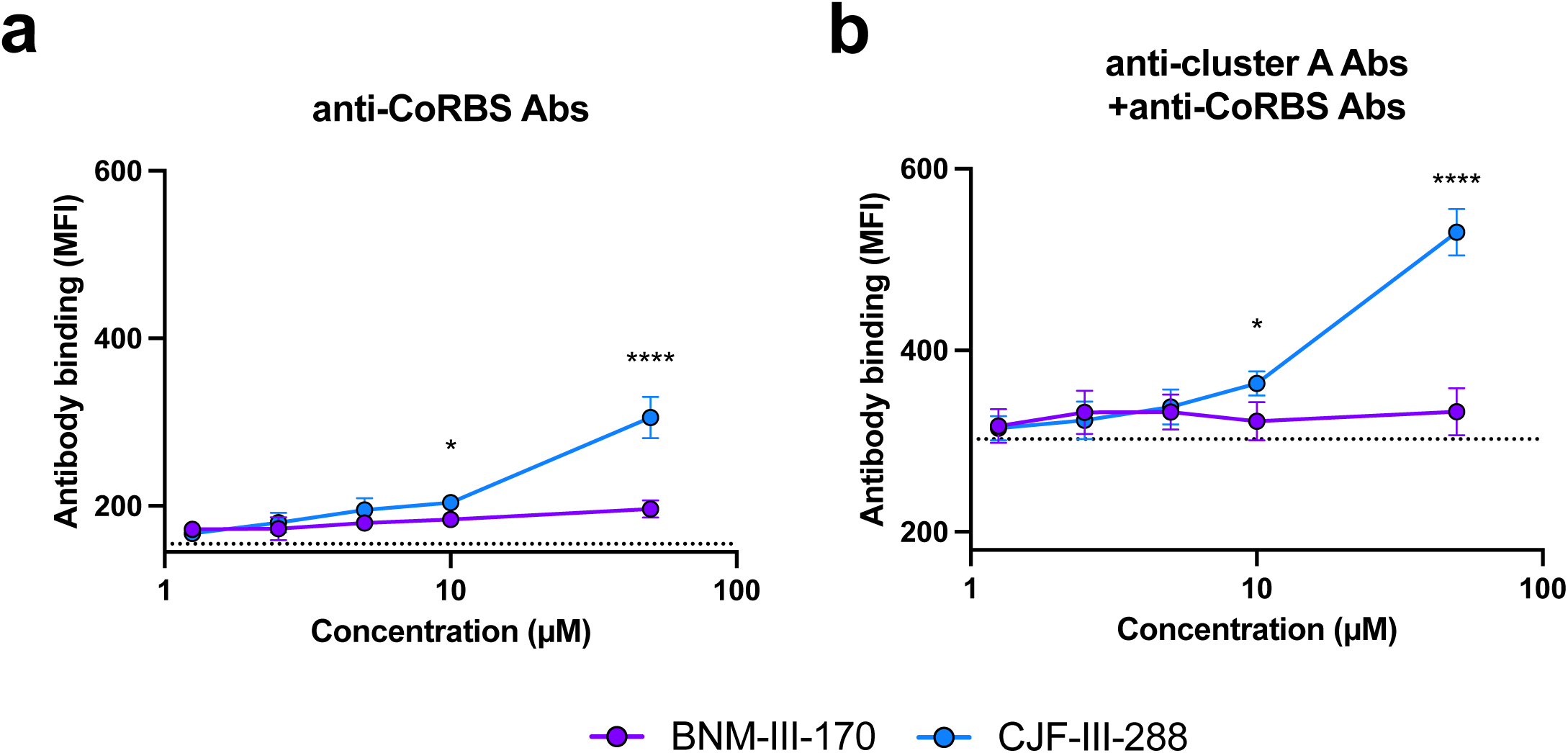
Improved recognition of HIV-1_BG505_-infected cells by nnAbs in presence of CJF-III-288. Recognition of HIV-1_BG505_-infected primary CD4+ T cells by anti-CoRBS Abs **a,** alone or **b,** in combination with anti-cluster A Abs, in the presence of indicated concentration of BNM-III-170 or CJF-III-288. Graphs represent the median fluorescence intensities (MFI) obtained in at least 4 independent experiments. The dashed lines represent the mean binding obtained in absence of CD4mc. Statistical significance was tested using Two-way ANOVA test with a Holm-Sidak post-test (* p<0.05,**** p<0.0001).

**Extended data Fig. 3.**
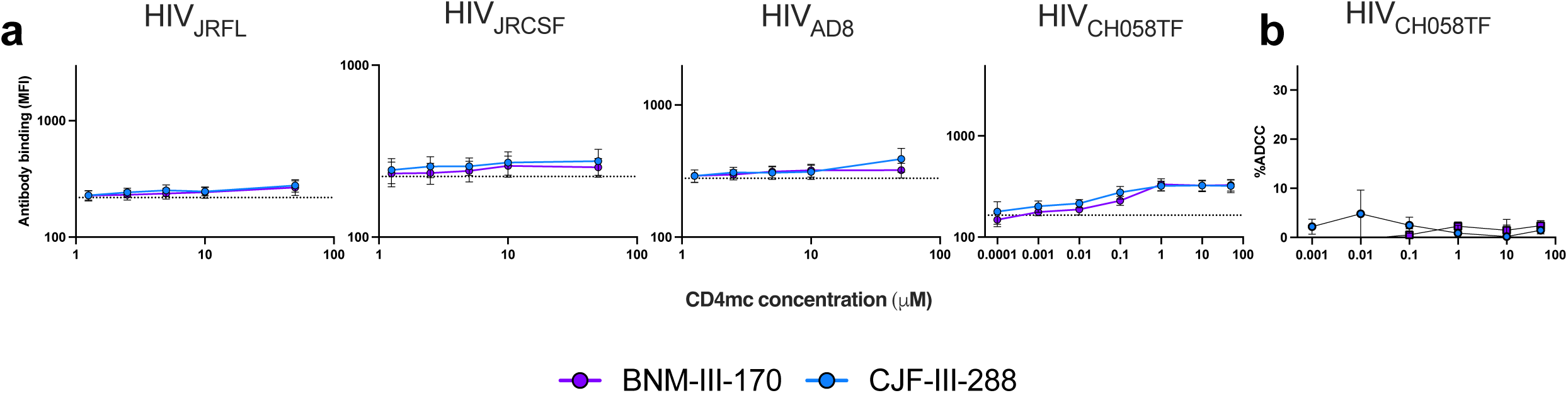
A32 binding and ADCC activity in presence of different concentrations of CD4mc. **a**, Recognition of primary CD4+ T cells infected with indicated IMC by the anti-cluster A Abs A32 in the presence of indicated concentration of BNM-III-170 or CJF-III-288. Graphs represent the median fluorescence intensities (MFI) obtained in at least 4 independent experiments with each IMCs. **b,** ADCC-mediated elimination of CD4+ T cells infected HIV-1_CH058TF_ by A32 in the presence of indicated concentration of BNM-III-170 or CJF-III-288. Shown are percentage of ADCC obtained in at least 4 experiments. The dashed lines represent the mean binding or ADCC obtained in absence of CD4mc.

**Extended data Fig. 4.**
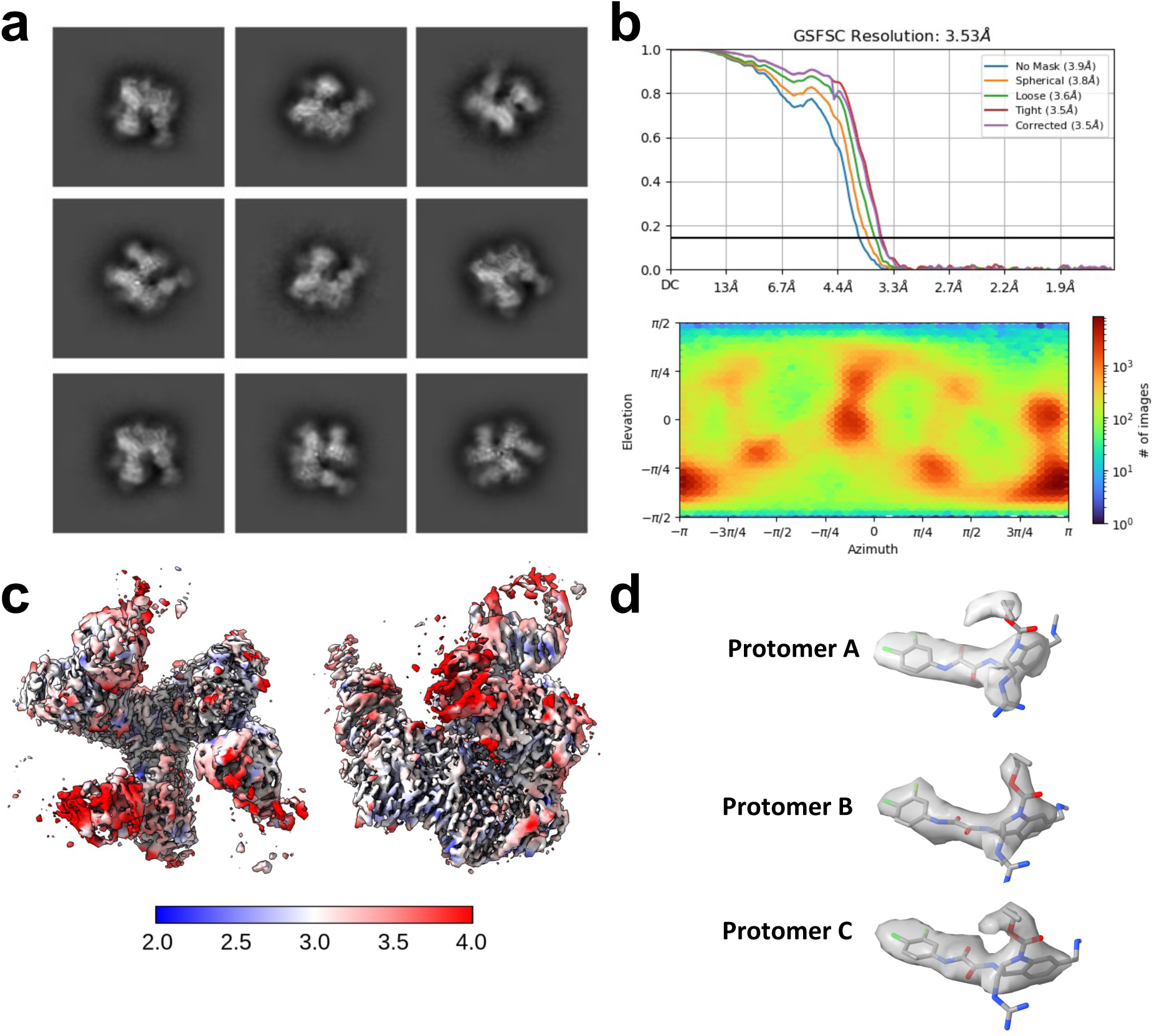
Cryo-EM structure of the CJF-III-288-BG505 SOSIP.664-17b Fab complex. **a**, Selected 2D classes for ab initio map reconstruction. Micrographs were collected using on a FEI Titan Krios electron microscope operating at 300 kV equipped with Gatan Bioquantum Image filter-K3 direct electron detector (Gatan Inc). **b**, The Fourier shell correlation curves (FSC cutoff 0.143) from the final non-uniform refinement step and the direction distribution plot of all particles used in the final refinement. **c**, Local resolution estimation. **d**, Density and corresponding model for CJF-III-288 from each protomer (protomer A corresponds to chain C, protomer B corresponds to chain A and protomer C corresponds to chain E).

**Extended data Fig. 5.**
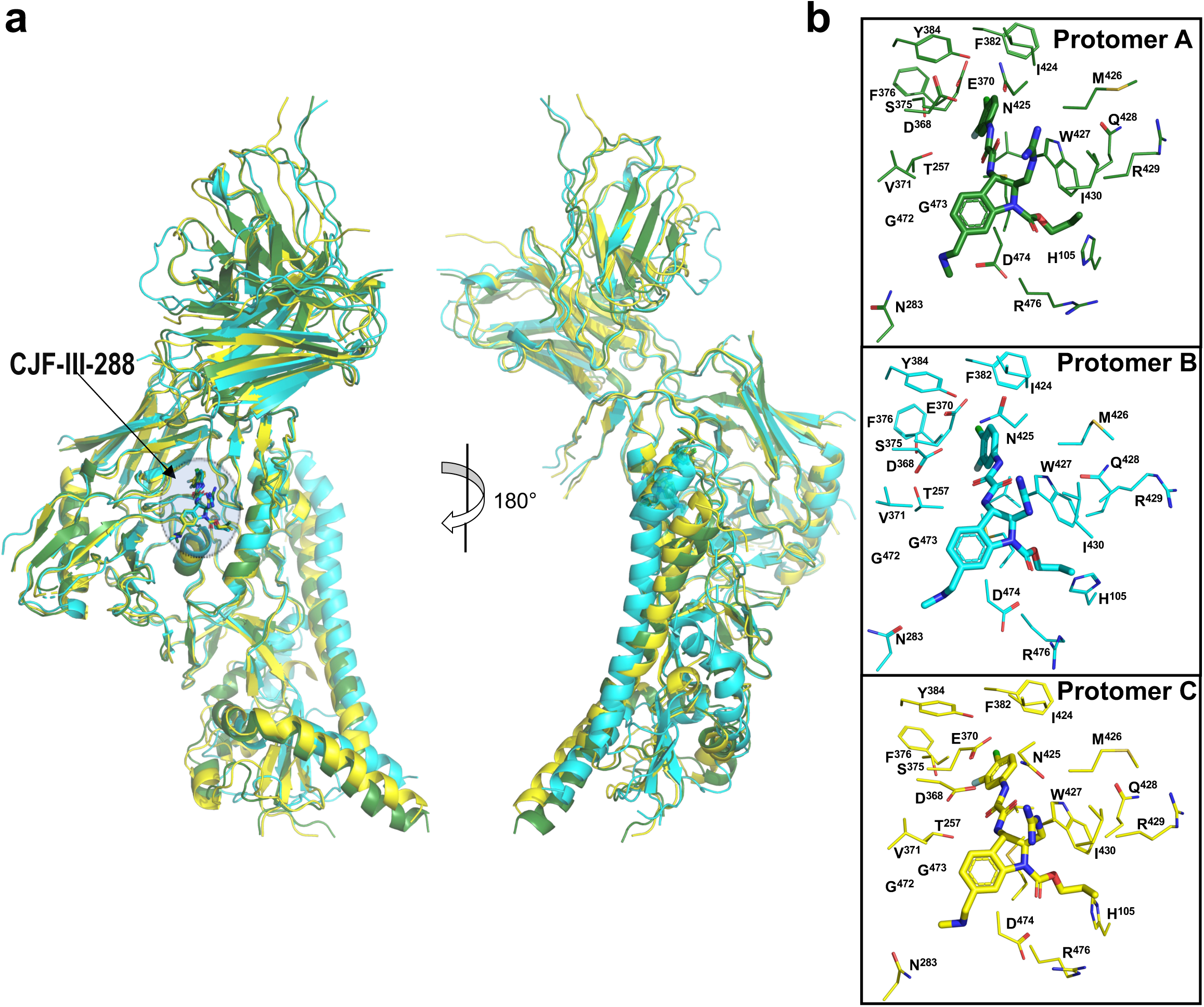
Superimposition of the three protomers of the asymmetric trimer of the CJF-III-288-BG505-17b Fab complex. **a**, Superimposition of protomers. 17b, gp120 and gp41 in each protomer are shown as cartoons and CJF-III-288 is shown as sticks. Protomer A is colored green, protomer B colored cyan and protomer C colored yellow. The protomers were superimposed and the root-mean-square deviation (RMSD) values of Cα atoms between protomers calculated. The RMSD between protomers A and B, protomers A and C, and protomers B and C are 2.7 Å, 1.73 Å, 3.14 Å, respectively. **b**, Residues forming the binding pockets in each protomer. The RMSD values of Cα atoms of pocket residues are 0.565 Å, 0.563 Å, 0.524 Å for protomers A and B, protomers A and C, and protomers B and C, respectively.

**Extended data Fig. 6.**
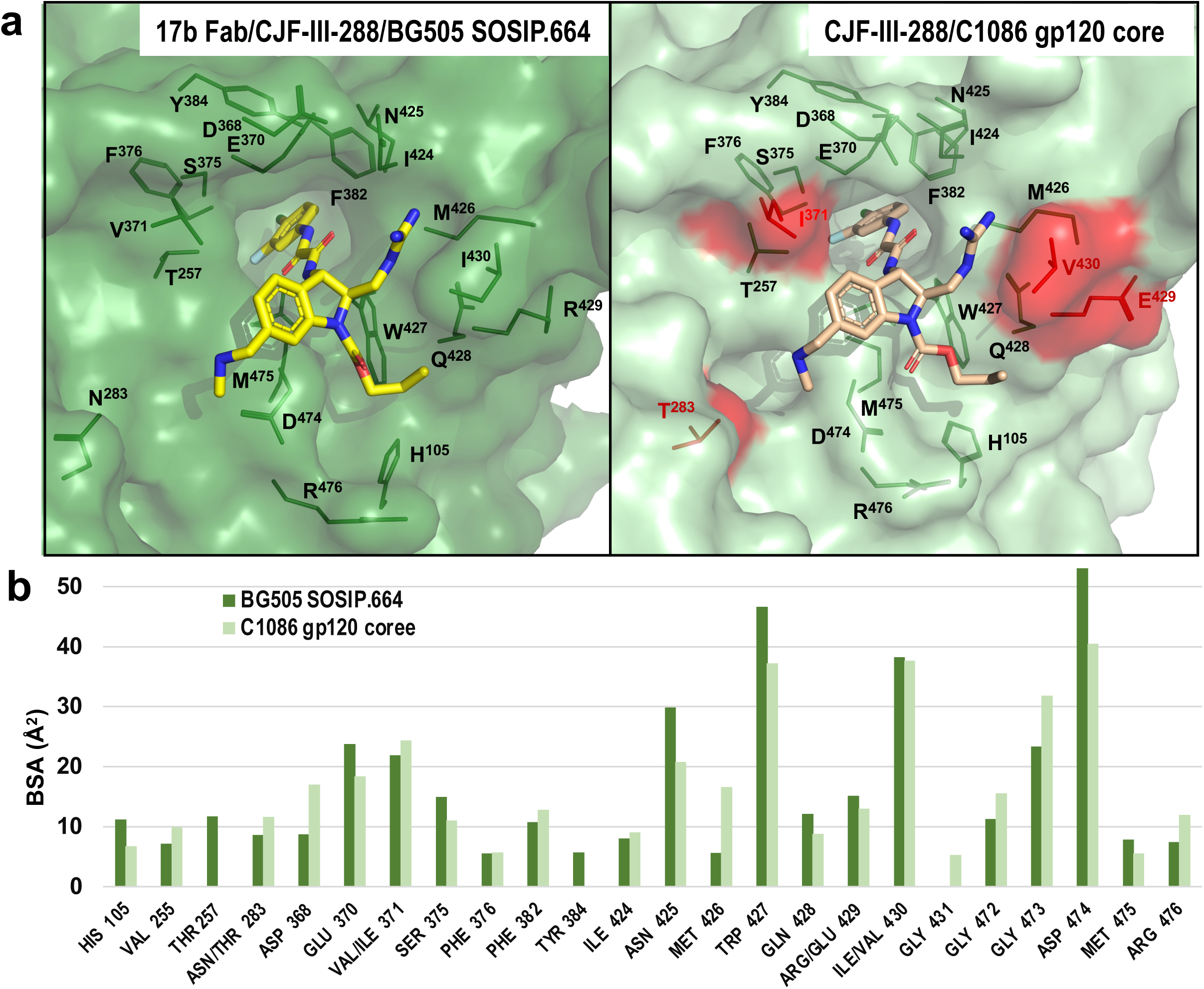
Comparison of the CJF-III-288 binding pockets as in the CJF-III-288-BG505-17b Fab and the CJF-III-288-C1086 gp120 core_e_ complexes. **a**, The CJF-III-170 binding pocket (from Protomer A) is superimposed based upon gp120 to the complex of the CJF-III-288 gp20 core_e_ (8FM3). Strain specific residues in CJF-III-288-gp120 core_e_ complex are highlighted red. **b**, Buried surface area (BSA) of gp120 residues buried at the CJF-III-288 binding interfaces.

**Extended data Fig. 7.**
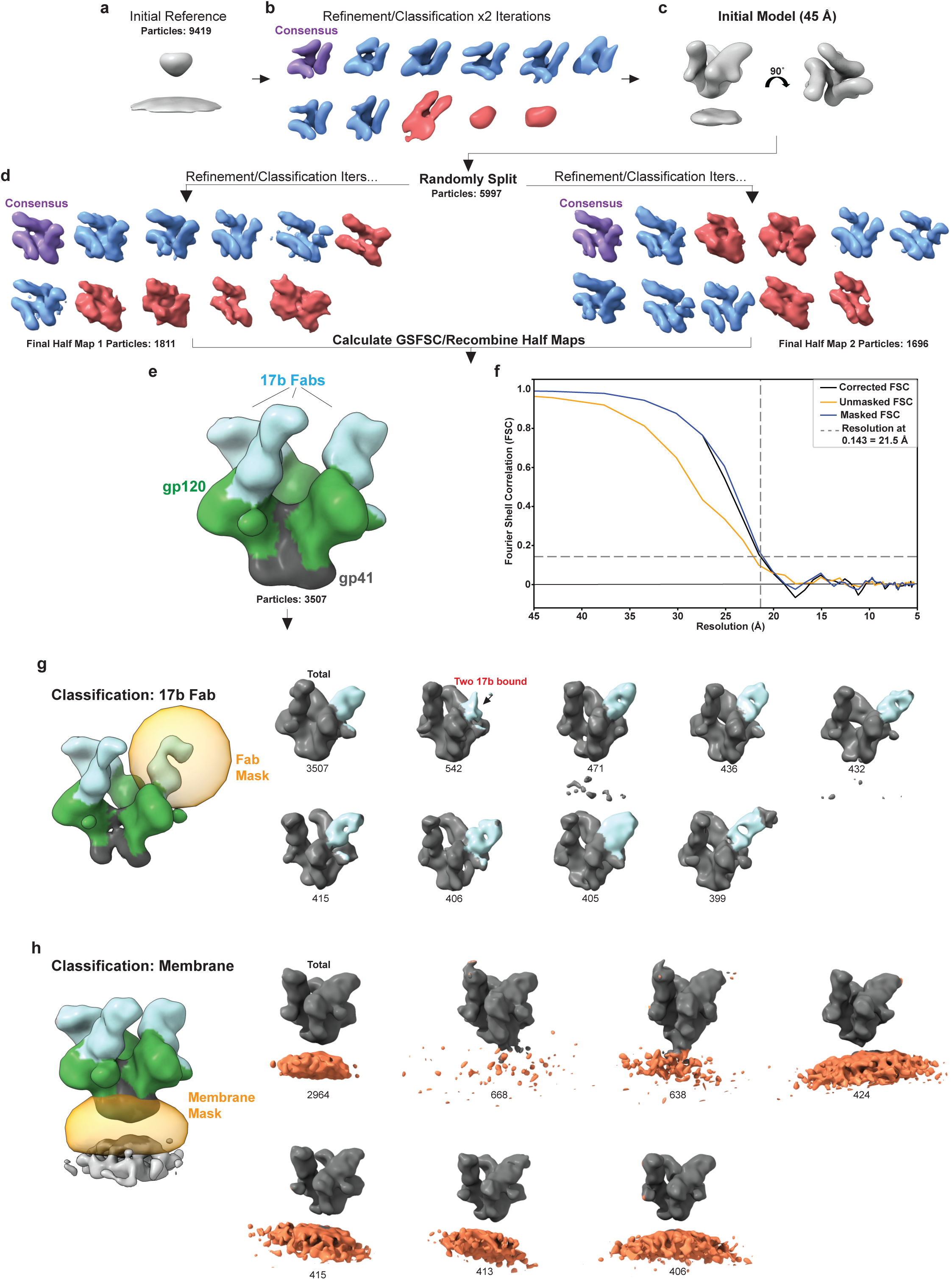
CryoET refinement and classification workflow. **a**, HIV-1_BaL_ Env trimers in complex with CJF-III-288 and 17b Abs, along with the center of each virion, were manually picked using IMOD software. PEET program spikeInit was used to determine approximate particle orientations to generate an initial reference. **b,c**, Refinement and classification were performed to remove junk particles (red) and generate an initial model. Resolution was limited to 45 Å by strong lowpass filtration and binning. **d,** Particles were randomly split into two halves for further refinement. Classification was performed to select Env trimers bound to three 17b Abs (blue). **e**, Isosurface rendering of the structure. **f**, Fourier shell correlation calculated from the two half-maps. Resolution was estimated at FSC = 0.143. **g**, Masked classification of the 17b Fab domain with weaker density shows heterogeneity in the 17b binding orientation. **h**, Masked classification on the membrane for Env bound to three 17b shows tilting on the membrane. The number of particles in each subclass for **g** and **h** are shown.

**Extended data Table 1.** Cryo-EM data collection and refinement statistics.

| **BG505 SOSIP.664 in complex with 17b Fab and CJF-III-288** |  |
| --- | --- |
| EMDB | EMD-45530 |
| PDB | 9CF5 |
| **Data Collection and Processing** |  |
| Microscope | FEI TITAN KRIOS G1 |
| Voltage (kV) | 300 |
| Detector | Gatan K3 |
| Energy filter (slit) | Yes (20 eV) |
| Cs corrector | No |
| Magnification | 105,000 x |
| Pixel Size (Å/px) | 0.832 (0.416 superres) |
| Total electron dose (e<sup>-</sup>/Å<sup>2</sup>) | 54.2 |
| Dose rate (e<sup>-</sup>/px/s) | 15 |
| Defocus Range (µm) | 0.5-2.7 |
| Micrograph collected | 9,431 |
| Number of picked particles | 6,768,076 |
| Box size (pix) | 400 |
| Number of final particles | 1,087,694 |
| Symmetry | C1 |
| Resolution (Å) (FSC 0.143) | 3.5 |
| B-factor for map (Å<sup>2</sup>) | 159.5 |
| **Refinement (Phenix) & validation** |  |
| Model composition |  |
| Non-hydrogen atoms | 18,048 |
| Protein | 17,083 |
| Ligands (CJF-III-288) | 111 |
| Average B-factors (Å<sup>2</sup>) |  |
| Protein | 101 |
| Ligands (CJF-III-288) | 54 |
| CC_mask | 0.738 |
| EMRinger Score | 1.04 |
| RMSD Bond lengths (Å) | 0.004 |
| RMSD Bond angles (°) | 0.685 |
| Molprobity score | 2.17 |
| Clash score | 15.2 |
| Rotamer outliers (%) | 0.53 |
| Ramachandran Plot |  |
| Favored (%) | 92.2 |
| Allowed (%) | 7.8 |
| Disallowed (%) | 0 |

